# Time Course and Microcircuit Mechanisms of Primary Motor Cortex Dysfunction in Progressive Parkinsonism

**DOI:** 10.64898/2026.07.27.741031

**Authors:** Liqiang Chen, Hiba Douja Chehade, Samhitha Somavarapu, Hong-Yuan Chu

## Abstract

Dysfunction of the primary motor cortex (M1) has long been implicated in the pathophysiology of Parkinson’s disease (PD), mostly as a link by which abnormal basal ganglia and thalamic activities are translated into motor symptoms. However, emerging evidence suggests that M1 neurons also exhibit maladaptive changes in advanced parkinsonism. Here, using a progressive mouse model of nigrostriatal neurodegeneration (i.e., the MitoPark mice, MP), we found that M1 neurons develop age- and striatal dopamine-dependent synaptic and cellular adaptations as parkinsonism progresses. The optogenetic, electrophysiological, pharmacological, and CRISPR-mediated genetic studies demonstrated that impaired α5-GABA_A_ receptor-mediated inhibition and excessive activation of NMDA receptors of M1 pyramidal neurons are key microcircuit mechanisms underlying cortical circuit remodeling during progressive striatal dopamine (DA) loss. Furthermore, we found that treatment with L-DOPA at early or late stages of parkinsonism can prevent or rescue, respectively, the synaptic and cellular adaptations in M1. Together, the present study demonstrates the time course and the underlying molecular and microcircuit mechanisms of cortical network dysfunction during the development of parkinsonism.

## Introduction

Parkinson’s disease (PD) is a progressive neurodegenerative disorder characterized by the gradual loss of dopaminergic neurons in the substantia nigra pars compacta (SNc) and their axonal projections to the striatum. Chronic striatal dopamine (DA) depletion alters the anatomical and physiological integrity of the basal ganglia^1,2^, resulting in pathological neural activity within the cortico-basal ganglia-thalamocortical network^1,3–5^. These network abnormalities underlie the traditional motor symptoms of PD (e.g., akinesia, bradykinesia, and rigidity). The abnormal neuronal activity manifests as altered firing frequency, an increased irregularity of firing, and synchronized burst firing across basal ganglia nuclei, including the substantia nigra pars reticulata (SNr)^1,5^. The pathological GABAergic output of the SNr is thought to propagate to the cerebral cortex via the ventral thalamus, potentially triggering (mal)adaptive plastic changes within the thalamocortical network that contribute to both motor and nonmotor symptoms^6,7^.

The primary motor cortex (M1) is a laminar structure comprising heterogeneous neuronal subpopulations. By forming complex microcircuits, distinct cortical neuron subtypes integrate sensory and cognitive information to generate motor commands essential for initiating and executing voluntary movement^8,9^. Consequently, functional impairment or neuronal loss in M1 contributes to numerous neurological disorders^6,10,11^. The traditional model of the pathophysiology of PD assigns M1 an important role in the emergence of parkinsonism^1,3,4^. Moreover, earlier work in MPTP-treated nonhuman primates (NHPs) reported that degeneration of the SNc DA neurons alters the firing frequency and pattern of M1 pyramidal neurons both at rest and during motor tasks^12–14^. Notably, layer 5 pyramidal tract (PT) neurons fired less and showed an enhanced tendency to discharge in bursts in parkinsonian monkeys, whereas corticostriatal projection neurons remained largely unaffected^13^. Aligning with these findings, our recent work using *in vivo* Ca^2+^ imaging from the MitoPark (MP) mice demonstrated that progressive loss of SNc DA neurons decreases the activity of PT neurons, but not that of intratelencephalic (IT) neurons that give rise to corticostriatal projections, in M1 during skilled movements^15^. Because the motor thalamus provides major glutamatergic synaptic excitation to M1, conveying basal ganglia output back to the cortex. Therefore, pathological inputs from the motor thalamus^16,17^ may drive the abnormal firing rates and patterns observed in PT neurons but not IT neurons in the parkinsonian state. However, this speculation is not supported by the equal innervation of PT and IT neurons by thalamic inputs under physiological state^18,19^, indicating that M1 dysfunction involves cortical microcircuit adaptations in parkinsonism.

At the cortical level, we recently demonstrated numerous synaptic and cellular alterations specific to PT neurons in mouse models of parkinsonism^19–21^. Specifically, we found that, following the loss of SNc DA neurons, M1 PT neurons exhibited decreased intrinsic excitability and reduced responses to excitatory inputs from the motor thalamus^19,20^. In contrast, the cellular excitability and thalamic excitation of IT neurons were not altered^19,20^. However, the temporal progression of these M1 circuit alterations during the gradual loss of SNc DA neurons, and the associated microcircuit mechanisms, remain undetermined. Answering these questions is essential for designing effective, stage-specific cortical interventions for PD therapy^6,22^.

In the present work, we employed a progressive mouse model of nigrostriatal neurodegeneration, the MitoPark mice^15,23–25^, complemented with the broadly used, subchronic model of advanced stages of parkinsonism, the 6-hydroxydopamine (6-OHDA)-lesioning mice, to answer the following questions: What is the time course of M1 circuit adaptations during the progressive striatal DA depletion? What are the key molecular and microcircuit mechanisms underlying M1 circuit changes during PD progression? Do M1 circuits respond to dopaminergic medications at distinct stages of parkinsonism, and what mechanism(s) underlie such responses? We report that M1 circuit changes are first detectable in the moderately severe parkinsonian state, associated with > 80% striatal DA depletion. We further demonstrate that impaired cortical GABAergic inhibition and subsequent excessive activation of GluN2B-containing NMDA receptors (NMDARs) trigger the cellular and synaptic adaptations in M1 during the gradual loss of SNc DA neurons. Using pharmacological and chemogenetic approaches, we also show that thalamocortical synaptic deficits can be rescued by reducing the basal ganglia output. These results defined the time course of cortical circuit adaptations during progressive loss of SNc DA neurons and reveal several underlying molecular and microcircuit mechanisms.

## Results

We used the MP mice (Table S1: Key resource table) to study the time course of synaptic and cellular adaptations in M1 circuits during the gradual loss of SNc DA neurons. MP mice developed an age-dependent degeneration of the nigrostriatal innervation, as indicated by a gradual decrease in tyrosine hydroxylase (TH)-immunoreactivity (ir) in the dorsal striatum (Figure S1A-F). In our hands, the levels of striatal TH-ir in 8-, 12-, 16-, and 24-week-old MP mice were 99%, 54%, 19%, and 10% of controls, respectively. Behaviorally, the MP mice exhibited progressive impairment of locomotor activity in an open-field arena (Figure S1G, H)^23^. These observations confirm that MP mice develop adult-onset, progressive degeneration of the nigrostriatal dopaminergic projections and associated parkinsonian motor deficits, as previously reported^15,23–27^. Based on the temporal profiles of the nigrostriatal DA degeneration and the emergence of motor deficits in MP mice (Figure S1), we performed a longitudinal investigation of the M1 circuits at the normal (8 weeks), early motor (12 weeks), moderate motor (16 weeks), and advanced motor stages (24 weeks) of parkinsonism.

## Time course of synaptic adaptations in M1 during progressive loss of SNc DA neurons

Proper motor control and motor learning depend on normal connectivity within the thalamocortical network^18,28–31^. However, following a nearly complete loss of SNc DA neurons, the synaptic strength of the thalamic input onto PT neurons (thalamo-PT) is markedly reduced^19^. To determine the time course of thalamo-PT synaptic adaptation during the progressive SNc DA neurodegeneration, we injected adeno-associated virus (AAV) encoding ChR2(H134R)-eYFP into the motor thalamus (i.e., ventromedial and ventrolateral thalamus) and Retrobeads into the pons to retrogradely label M1 PT neurons in MP mice and littermate control animals of various ages (Figure 1A). Three weeks post-surgery, we performed *ex vivo* whole-cell voltage-clamp recordings of optogenetically-evoked monosynaptic excitatory postsynaptic currents (oEPSCs, Figure 1B) from labeled PT neurons using an extracellular solution containing SR95531(GABAzine)/tetrodotoxin (TTX)/4-aminopyridine (4-AP)^19,32^. We measured the amplitude of oEPSCs at -80 mV to quantify the AMPA receptor (AMPAR)-mediated thalamo-PT synaptic strength during progressive loss of the SNc DA neurons (Figure 1C-F). At 8 and 12 weeks of age, there was no difference in the amplitude of thalamo-PT oEPSCs in response to different intensities of light stimulation between MP mice and littermate controls (Figure 1B-D). In moderately parkinsonian animals (16 weeks of age) and more severely affected animals (24 weeks), MP mice exhibited a significant reduction in the amplitude of thalamo-PT oEPSCs relative to their respective littermate controls (Figure 1B, E, F). As reported in the 6-OHDA-lesioned mice^19^, there was a significant reduction in the frequency of Sr^2+^-induced, optogenetically-evoked asynchronous miniature EPSCs at the thalamo-PT synapses in 24-week-old MP mice relative to littermate controls (Figure S2A-C). This indicates decreased numbers of functional connections between the motor thalamus and PT neurons in M1. There was no change in the paired pulse ratio or AMPA/NMDA ratio at the thalamo-PT synapses between MP mice and littermates at 24 weeks of age (Figure S2D-E), as seen in 6-OHDA-lesioned mice^19^. To determine whether the alterations in M1 were cell-subtype-specific, we also studied the thalamic inputs to IT neurons in layer 5 of M1 (i.e., “thalamo-IT” synapses). There were no differences in the amplitudes of thalamo- IT EPSC between MP mice and littermates at 24 weeks of age (Figure S3A, B). This observation suggests that thalamic input to IT neurons is not altered in MP mice, similar to findings in 6-OHDA-lesioned mice^19^. Taken together, we conclude that impairments of the thalamo-PT synaptic connectivity are induced at moderate motor stages of parkinsonism associated with >80% loss of striatal DA^25^.

**Figure 1.**
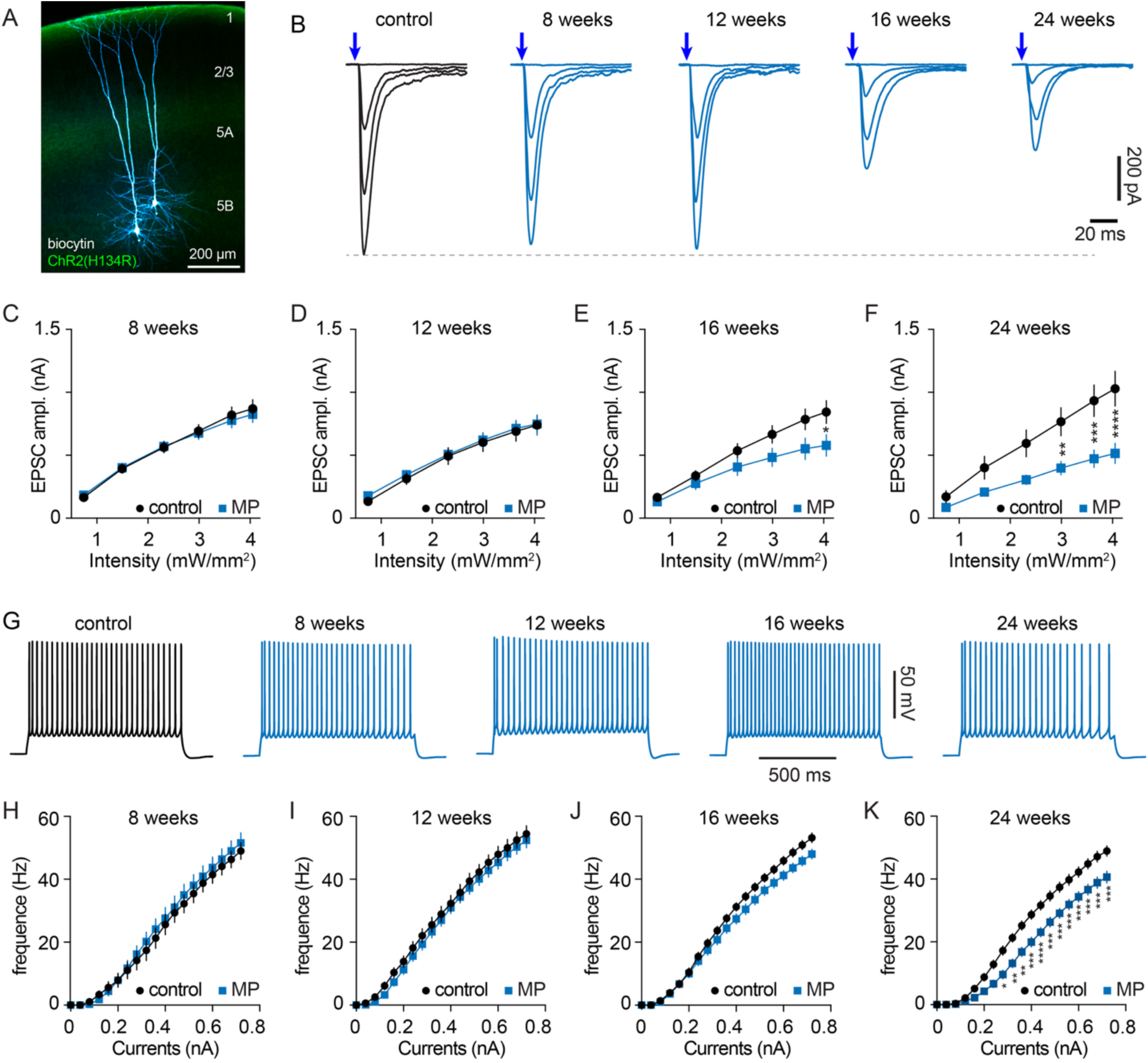
Time course of synaptic and cellular plasticity of M1 during progressive parkinsonism. **A)** Representative biocytin-labeled PT neurons in M1 (blue) and ChR2-expressing axon terminals (green) of the motor thalamic neurons. **B**) Representative EPSC traces from M1 PT neurons evoked by optogenetic stimulation of the thalamic axon terminals of control mice and MP mice across ages. EPSCs were evoked by delivering 1 ms light pulses at stepwise increasing intensities (0, 0.8, 2.3, 4.0 mW/mm^2^). Blue arrows indicate light pulses. **C-F**) Summary graphs showing progressive reduction of the amplitude of the thalamo-PT EPSCs in MP mice relative to littermate controls at 8- (36 cell/ 5 control mice and 24 cells/3 MP mice), 12- (42 cells/ 6 control mice and 48 cells/7 MP mice), 16- (35 cells/ 8 control mice and 35 cells/8 MP mice) and 24- (18 cells/ 4 control mice and 24 cells/5 MP mice) weeks of age. Blue arrows indicate light pulses. **G**) Representative AP traces of PT neurons evoked by somatic current injection (320 pA, 1 sec) from controls and MP mice across ages. **H-K**) Frequency-current (F-I) curves of PT neurons from MP mice and littermate controls at 8- (24 cells/ 6 control mice and 19 cells/5 MP mice), 12- (41 cells/6 control mice and 43 cells/6 MP mice), 16- (47 cells/10 control mice and 48 cells/10 MP mice) and 24- (52 cells/ 10 control mice and 45 cells/11 MP mice) weeks of age. C-F, H-K: ns, not significant, *p<0.05, **p < 0.01, ***p < 0.001, ****p < 0.0001, two-way repeated measures (RM) ANOVA followed by Sidak tests.

## Time course of intrinsic adaptations in M1 during progressive loss of SNc DA neurons

Previous studies had also demonstrated dampened intrinsic excitability in M1 PT neurons under parkinsonian conditions^19,20^. Since synaptic and intrinsic plasticity can be mediated by distinct molecular mechanisms^33,34^, we next determined the temporal relationship between the striatal DA depletion and intrinsic adaptations in M1 in the parkinsonian state. With the blockade of ionotropic glutamatergic and GABAergic receptors by a cocktail of 6,7-dinitroquinoxaline-2,3-dione (DNQX,10 μM), D-2-amino-5-phosphonovaleric acid (D-APV, 50 μM), and GABAzine (10 μM), retrogradely labeled PT neurons were recorded and depolarized by a family of 1-second somatic current injections under the whole-cell current-clamp mode. The number of action potentials generated by the current injection was used as a measure of intrinsic excitability of the neurons. At 8 and 12 weeks of age, the intrinsic excitability of PT neurons did not differ between MP mice and littermate controls (Figure 1G-I). At 16 weeks, although there was a trend of reduced intrinsic excitability of PT neurons from MP mice compared to littermate controls, such differences did not reach statistical significance (Figure 1J). At 24 weeks of age, MP mice showed a significant reduction in the intrinsic excitability of PT neurons relative to littermate controls (Figure 1K). These data demonstrate the temporal trajectory of intrinsic adaptations in PT neurons that develop over the course of gradual striatal DA depletion. Importantly, there was no change in the intrinsic excitability of M1 IT neurons from 24-week-old MP mice relative to littermate controls (Figure S3C, D), confirming that the parkinsonism-related alterations in intrinsic excitability of M1 neurons are cell-type-specific^20^.

In our previous experiments in 6-OHDA-lesioned mice, we found that the impaired intrinsic excitability of PT neurons was mainly mediated by dysfunctional persistent Na^+^ channels, resulting in a depolarized action potential (AP) threshold and robust spike adaptation during prolonged membrane depolarization^20^. Consistently, PT neurons from MP mice at 24 weeks of age showed stronger spike adaptation and a more depolarized AP threshold than those from littermate controls (Figure S4). Therefore, we posit that the ionic mechanisms underlying the impaired PT neuronal excitability are partially shared between MP mice with advanced parkinsonism and 6-OHDA-lesioned mice.

Together, the data above suggest that robust changes in intrinsic excitability of PT neurons are induced at an advanced stage of parkinsonism associated with nearly complete loss of striatal DA (i.e., < 10% remaining)^25^.

### Impaired cortical GABAergic inhibition during progressive loss of SNc DA neurons

The results described above presented the time course of synaptic and intrinsic plasticity within the thalamocortical circuits during the progressive loss of SNc DA neurons. We next aimed to determine the key molecular mechanisms underlying the observed plasticity in M1 circuits. Previous studies implicated that NMDARs may mediate synaptic and cellular deficits of PT neurons in parkinsonian conditions^19^. It has been shown that a disrupted interaction between GABAergic inhibition and NMDAR-mediated excitation critically contributes to basal ganglia circuit dysfunction in parkinsonism^35^. Thus, we tested the hypothesis that impaired GABAergic signaling promotes NMDAR activation, triggering cellular and synaptic adaptations in M1 PT neurons following nigrostriatal DA degeneration.

To measure overall GABAergic inhibition, we first recorded spontaneous inhibitory synaptic currents (sIPSCs) from retrogradely labeled PT and IT neurons in 6-OHDA-lesioned mice and vehicle-injected controls using a high Cl^-^ internal solution. We found that the sIPSC frequency of PT neurons decreased significantly in 6-OHDA-lesioned mice relative to vehicle-injected controls (Figure S5A, C). In striking contrast, there was no difference in the sIPSC frequency recorded in IT neurons between 6-OHDA-lesioned mice and vehicle-injected controls (Figure S5B, C). Moreover, the sIPSC frequency was significantly higher in PT neurons than in IT neurons from vehicle-injected controls (Figure S5A-C), suggesting that PT neurons are under more potent GABAergic inhibition than IT neurons. These data suggest that GABAergic interneurons exert cell-type-selective inhibition of PT and IT neurons in the healthy state and that loss of SNc DA neurons decreases GABAergic inhibition of PT neurons, but not that of IT neurons.

To determine the time course of impaired inhibition, we next examined GABAergic inhibition of PT neurons in MP mice of different ages. As predicted, the frequency of sIPSCs in PT neurons was significantly lower in 24-week-old MP mice relative to littermates (Figure S5E-F). Interestingly, PT neurons from MP mice at 16 weeks of age also showed a lower sIPSC frequency than controls (Figure S5D, F). These data suggest that PT neurons receive more robust GABAergic inhibition than IT neurons in the healthy state, and that impairments of cortical GABAergic inhibition begin at a moderate motor stage and persist as SNc DA neurons progressively degenerate.

## Asymmetric inhibition of PT and IT neurons by SST-INs

Next, we investigated interneuron subtypes that contribute to the asymmetric inhibition of PT and IT neurons. It has been reported that PT and IT neurons receive comparable perisomatic GABAergic inhibition from PV-INs, which is not altered following striatal DA depletion^21,36,37^. We therefore focused on the question of whether the dendrite-targeting SST-INs inhibit PT neurons more strongly than IT neurons. SST-INs innervate pyramidal neurons in the hippocampus and cortex through the α5-subunit-containing GABARs with slow kinetics^38,39^. First, we recorded sIPSCs from PT and IT neurons using a high Cl^-^internal solution with the blockade of fast glutamatergic transmission using DNQX (10 μM) and D-APV (50 μM), and tested the effects of an α5-GABAR inverse agonist (α5IA)^40^ on GABAergic inhibition in PT and IT neurons of control mice (Figure 2A-B). We found that bath application of α5IA (100 nM) significantly decreased the amplitude of sIPSCs in PT neurons, but not in IT neurons (Figure 2C), suggesting that α5-GABARs contribute differently to GABAergic inhibition of PT and IT neurons. Considering the unique biophysical properties of α5-GABARs, we attempted to isolate α5-GABA_A_R-mediated IPSCs based on their slow kinetics. We found that the distribution of the half-width of sIPSCs from PT neurons showed two peaks (Figure 2D), supporting the presence of fast (mean_1_ = 3.6 ms) and slow (mean_2_ = 10.3 ms) sIPSCs. In contrast, the sIPSC half-width in IT neurons showed a normal distribution with a single peak centered at 4.8 ms (Figure 2E), indicating a pronounced proportion of fast sIPSCs. In the following analysis, we chose a half-width cutoff of 8 ms to separate “slow” and “fast” sIPSCs. As expected, application of α5IA (100 nM) effectively reduced the amplitude of slow sIPSCs but had little effect on the amplitude of fast sIPSCs of PT neurons (Figure 2F). These data support that M1 PT and IT neurons receive distinct levels of α5-GABAR-mediated inhibition.

**Figure 2.**
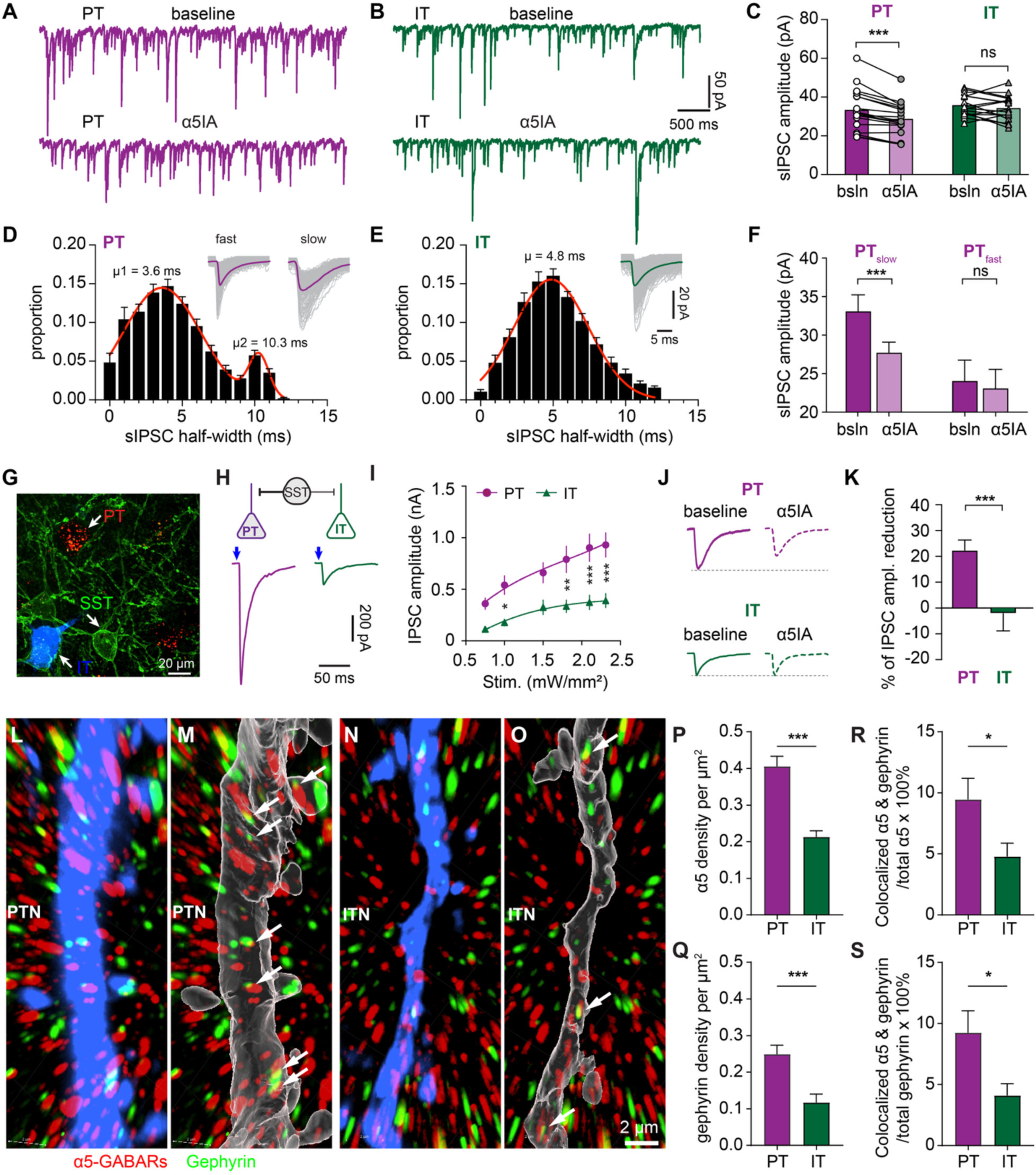
SST-INs exert asymmetric inhibition of PT and IT neurons via α5-GABA_A_Rs. A-B) Representative traces of sIPSCs recorded from PT (A) and IT (B) neurons before and after bath application of α5IA. sIPSC amplitude in PT neurons, baseline = 33.3±2.5 pA, n = 18 neurons/3 mice, α5IA = 28.6±2.0 pA, n = 18 neurons/3 mice, p < 0.0001; sIPSC amplitude in IT neurons, baseline = 35.7±1.3 pA, n = 17 neurons/3 mice, α5IA= 34.3±1.5 pA, n = 17 neurons/3 mice, p = 0.2; WSR test. **C**) α5IA reduced the amplitude of sIPSCs in PT neurons, but not IT neurons. **D-E**) Histogram showing the distribution of sIPSC amplitude in PT neurons (D) and IT neurons (E). Inset, the overlaid individual sIPSCs (gray) and the averaged traces (solid lines). **F**) α5IA reduced the amplitude of slow sIPSCs in PT neurons but had little effect on fast sIPSCs. IPSC_slow_, baseline = 33.3±2.2 pA, α5IA = 27.9±1.4 pA, n = 18 neurons/3 mice, p < 0.0001, WSR test; IPSC_fast_, baseline = 24.3±2.7 pA; α5IA= 23.3±2.5 pA, n = 18 neurons/3 mice, p = 0.15, WSR test; **G**) Representative confocal image showing the retrogradely labeled PT (red) and IT (blue) neurons in a brain slice of an SST-ChR2 mouse (green). **H**) Representative traces of oIPSC in PT and IT neurons. Blue arrows indicate light pulses. **I**) Summary graph showing the amplitude of oIPSCs in PT and IT neurons. PT: 23 neurons/5 mice, IT: 20 neurons/5 mice, two-way RM ANOVA followed by Sidak test. **J-K**) Effect of α5IA on the amplitude of oIPSCs in PT and IT neurons. %oIPSC reduction, PT = 22.4±3.9%, n = 12 neurons/3 mice, IT = -2.1±6.7%, n = 10 neurons/3 mice, p = 0.0015, MWU test. **L-O**) Representative confocal images of spines on apical dendrites of PT (L) and IT (N) neurons and the reconstructed masks (M, O). White arrows show examples of α5-GABAR puncta colocalized with gephyrin puncta. **P**) Bar graph showing differences in α5-GABARs between PT and IT neurons (PT = 0.41±0.03 puncta/µm^2^, n = 26 segments/5 mice, IT = 0.21±0.02 puncta/µm^2^, n= 21 segments/3 mice, p < 0.0001, MWU test). **Q**) Bar graph showing differences in gephyrin expression between PT and IT neurons (PT = 0.25±0.02 puncta/µm^2^, n = 26 segments/5 mice; IT = 0.12±0.02 puncta/µm^2^, n= 21 segments/3 mice, p < 0.0001, MWU test). **R**) Bar graph showing differences in percentage of colocalized α5+gephyrin puncta out of total α5 puncta (PT = 9.5±1.7%, n = 26 segments/5 mice; IT = 4.8±1.1%, n= 21 segments/3 mice, p = 0.0496, MWU test). **S**) Bar graph showing differences in percentage of colocalized α5+gephyrin puncta out of total gephyrin puncta (PT = 9.3±1.8%, n = 26 segments/5 mice; IT = 4.1±0.9%, n= 21 segments/3 mice, p = 0.04, MWU test). *p < 0.05, **p<0.01, ***p<0.001

We used the SST-ChR2-eYFP mice to directly quantify the synaptic strength of SST-IN inputs to PT and IT neurons (i.e., SST-PT and SST-IT synapses, respectively) and the involvement of α5-GABARs. To do this, we retrogradely labeled PT and IT neurons by injecting red Retrobeads and fast blue into the pons and contralateral striatum, respectively (Figure 2G). In whole-cell voltage-clamp mode, we recorded the optogenetically-evoked IPSCs (oIPSCs) from PT and IT neurons sequentially in the same slice (Figure 2G-H). We found that the amplitude of oIPSCs from PT neurons was greater than that from IT neurons (Figure 2H-I), confirming the asymmetric inhibition of PT and IT neurons by SST-INs. Application of α5IA (100 nM) significantly decreased the amplitude of SST-PT oIPSCs by 22.4%, but it produced little effect on the amplitude of SST-IT oIPSCs (Figure 2J-K). oIPSCs from both synapses were completely abolished by the selective GABA_A_R antagonist SR-95531 (GABAzine,10 μM). Although α5IA is selective for α5-GABARs, its maximal efficacy is only 30-40%^40^. After factoring in the intrinsic efficacy of α5IA, α5-GABAR activation accounts for 56-67% of the total IPSC amplitude at the SST-PT synapses. Thus, SST-INs differentially innervate PT and IT neurons, primarily via α5-GABA_A_Rs.

We also conducted super-resolution confocal imaging studies to visualize α5 subunits in the apical dendrites of retrogradely labeled and biocytin-filled PT and IT neurons (see Materials and Methods). To selectively study the synaptic α5 GABA_A_Rs, we quantified colocalization of α5 subunits with gephyrin, a postsynaptic scaffolding protein at GABAergic synapses. We found significantly higher densities of α5 subunits (Figure 2P) and gephyrin (Figure 2Q) in the dendrites of PT neurons than those of IT neurons. These data are consistent with the differences in functional GABAergic inhibition between PT and IT neurons (Figure 2A-C). Since α5 subunits locate both synaptically and extrasynaptically, we further determined whether synaptic α5 subunits differ between the two cell types. Indeed, there was a significantly higher percentage of colocalized α5 and gephyrin puncta out of total α5 or gephyrin immunoreactivity in PT neurons than in IT neurons (Figure 2R, S). Since gephyrin is a postsynaptic scaffold protein, these data suggest a higher proportion of α5 subunits are located within synaptic loci of PT neurons. Together, we conclude that SST-INs exert stronger α-5GABAR-mediated inhibition on PT than IT neurons.

## Impaired α5-GABARs function during progressive loss of SNc DA neurons

We next sought to determine whether the GABAergic inhibition of PT neurons by SST-INs (“SST-PT” synapses) is impaired in parkinsonism. MP and the acute 6-OHDA-lesioned mice shared many features of thalamocortical network dysfunction (Figure 1)^19,20^. It is likely that key neural mechanisms driving thalamocortical circuit dysfunction are conserved between the two models. Thus, we used these complementary models to study the microcircuitry basis and time course of M1 dysfunction in parkinsonism.

We injected 6-OHDA or vehicle into the MFB of SST-ChR2-eYFP mice to assess the strength of SST-PT synapses following a nearly complete loss of SNc DA neurons. In subsequent optogenetic experiments, we optically stimulated ChR2-expressing axon terminals in the superficial layers (L1-3, Figure 3A) to evoke SST-PT oIPSCs arising from the apical dendrites^41^. Using a high Cl^-^_ internal solution, we found that the amplitude of SST-PT oIPSCs was significantly reduced across all optical stimulation intensities in 6-OHDA-lesioned mice relative to vehicle-injected controls (Figure 3B, C), indicating that the SST-PT GABAergic inhibition is weakened after loss of SNc DA neurons. Furthermore, SST-PT synapses in vehicle-injected controls showed robust short-term depression during repeated optogenetic stimulation (5 pulses at 20 Hz, 1 ms in duration, Figure 3D-E). In contrast, this short-term depression was significantly attenuated in 6-OHDA-lesioned mice (Figure 3D-E), indicating a possible reduction in presynaptic release probability at the SST-PT synapses in parkinsonism. In vehicle-injected control mice, bath application of α5ΙΑ (100 nM) significantly decreased the amplitude of SST-PT oIPSCs, confirming the involvement of α5ΙΑ into SST-PT synaptic transmission (Figure 3F, G). In contrast, the effect of α5ΙΑ to SST-PT oIPSCs was absent in 6-OHDA-lesioned mice (Figure 3F, G). The results suggest that the synaptic strength of α5-GABAR-mediated SST-PT inhibition decreased following the loss of SNc DA neurons, thereby preventing further suppression by α5IA.

**Figure 3.**
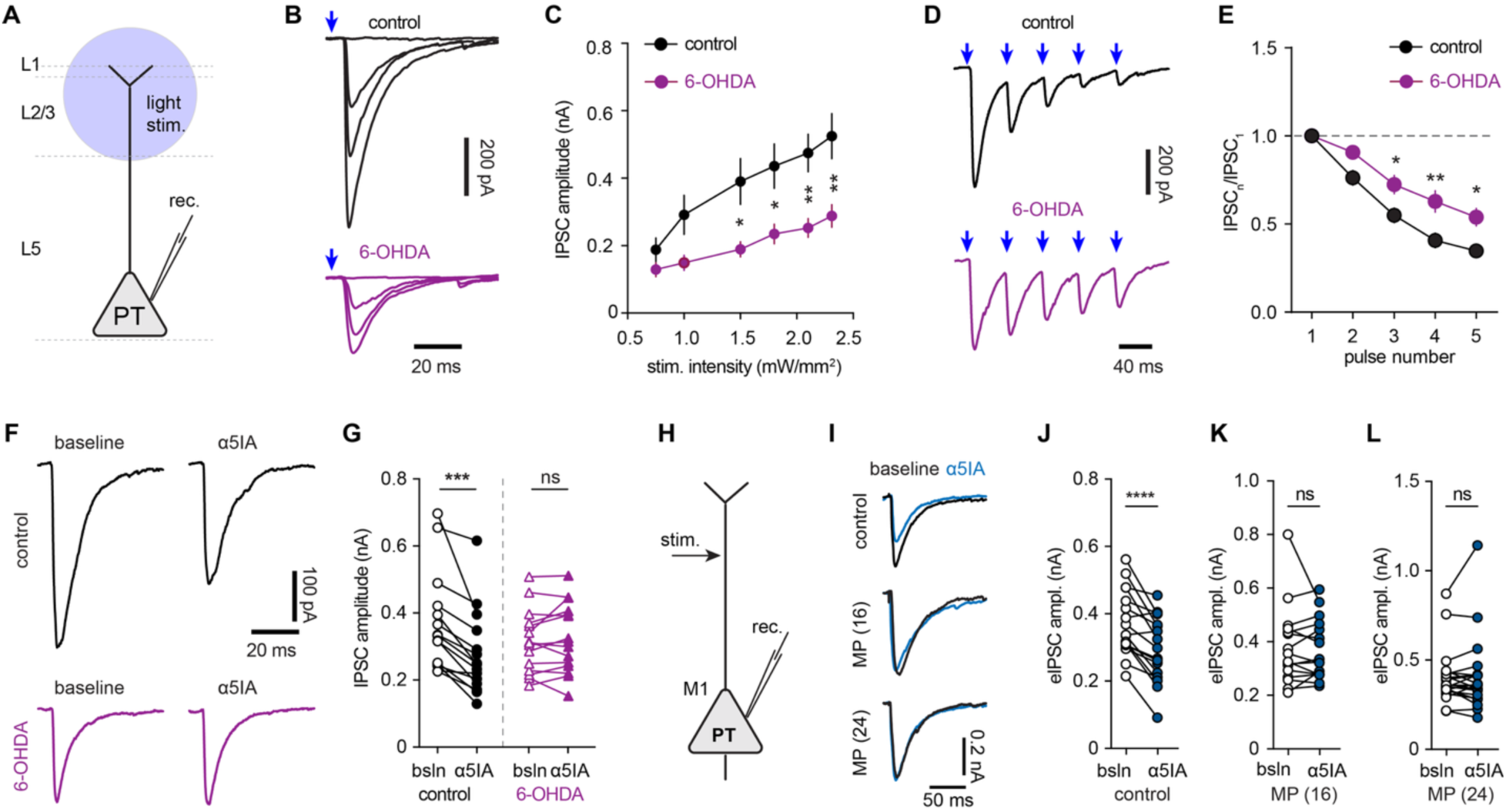
Impaired α5-GABA_A_R-mediated inhibition of PT neurons following the loss of SNc DA neurons. **A)** Experimental design. **B-C**) Representative traces (B) and the summarized results (C) showing oIPSCs arising from layers 1-3 of M1 between control and 6-OHDA-lesioned mice. Control = 18 neurons/4 mice, 6-OHDA = 16 neurons/4 mice, two-way RM ANOVA followed by Sidak test. **D-E**) SST-PT synapses show altered short-term plasticity in 6-OHDA-lesioned mice relative to controls. A train of oIPSCs was evoked by 5 pulses (1 ms in duration) of stimulation at 20 Hz. **F-G**) Effect of α5IA on the amplitude of oIPSCs in PT neurons from control and 6-OHDA-lesioned mice. Controls: baseline = 0.37±0.04 nA, α5IA= 0.28±0.03 nA, n = 15 neurons/4 mice, p < 0.0001; 6-OHDA mice: baseline = 0.32±0.024 nA, α5IA= 0.33±0.027 nA, n = 15 neurons/4 mice, p = 0.49; WSR test. **H**) Diagram of experimental design in MP mice. **I**) Representative eIPSC traces at baseline and after α5IA application from control and MP mice at 16- and 24-weeks of age. **J-L**) Summarized graphs showing the effect of α5IA on electrically-evoked IPSCs in PT neurons from controls (baseline = 0.37±0.02 nA, α5IA= 0.30±0.02 nA, n = 17 neurons/4 mice, p < 0.0001, WSR test), 16-week-old (baseline = 0.38±0.037 nA; α5IA= 0.37±0.028 nA, n = 16 neurons/4 mice, p = 0.94, WSR test), and 24-week-old (baseline = 0.40±0.034 nA, α5IA= 0.41±0.045 nA, n = 21 neurons/4 mice, p = 0.84, WSR test) MP mice. ns, not significant, *p< 0.05, **p < 0.01, ***p < 0.001.

To determine the time course of functional downregulation of α5-GABARs during progressive striatal DA depletion, we studied SST-PT synaptic properties in MP mice of different ages. Given that SST-INs preferentially target the dendrites of cortical pyramidal neurons, we electrically stimulated the L2/3 of M1 to evoke IPSCs arising from the apical dendrites of Retrobeads-labeled PT neurons (Figure 3H). The choice of electrical stimulation was based on the limited genetic tools available to target SST-INs in MP mice. We found that the amplitude of electrically-evoked IPSCs (eIPSCs) of retrogradely labeled PT neurons from littermate controls was significantly and consistently reduced by 100 nM α5IA (Figure 3I, J), indicating that the electrical stimulation approach recruited the dendrite-targeting SST-INs and the α5-GABA_A_R-mediated transmission. However, the effect of α5IA on the amplitude of eIPSCs in PT neurons was absent in the 16- and 24-week-old MP mice (Figure 3K, L). These data suggest that progressive loss of DA persistently weakens α5-GABAR-mediated inhibition of PT neurons by SST-INs, being identifiable at a moderate motor stage of parkinsonism with ∼80% striatal DA depletion.

## Impaired SST-IN inhibition promotes NMDAR activation at the thalamo-PT synapses during progressive loss of SN DA neurons

Thalamic inputs primarily target dendritic spines of M1 pyramidal neurons, which are concurrently innervated by the GABAergic axon terminals^42,43^. SST-INs are the dendrite-targeting interneurons that produce prolonged inhibition of pyramidal neurons via slow kinetic α5-GABARs to modulate NMDAR-mediated dendritic excitation^38^. Our recent work and others reported that NMDARs are enriched at the thalamo-PT synapses and their activation likely contribute to the decreased AMPAR-mediated neurotransmission in parkinsonism^19,20,30^. Yet, when and how NMDARs become overactive during progressive DA depletion remains to be determined. We determined whether impaired α5-GABAR inhibition and NMDAR overactivation at thalamo-PT synapses are linked during progressive striatal DA depletion, so that the impaired function of α5-GABARs promotes the persistent activation of NMDARs, as in the hippocampal circuits^38^.

We injected the AAV9-hSyn-ChR2(H134R) into the motor thalamus and Retrobeads into the pons of the MP mice and littermate controls to assess α5-GABAR- mediated modulation of NMDAR activation at the thalamo-PT synapse (Figure 4A). Notably, application of α5IA slightly depolarized the resting membrane potential (RMP) of neurons in both control and MP mice. All subthreshold and suprathreshold responses were recorded at RMPs by injecting a small holding current to compensate for this shift. The holding currents injected were comparable between controls and MP mice. At the RMP, optogenetic stimulation of ChR2-labeled axons evoked robust postsynaptic potentials (PSPs) in retrogradely labeled PT neurons of control mice (Figure 4B, black). Application of the α5IA (100 nM) significantly increased the integral of PSPs (i.e., area under the curve) evoked at RMP (Figure 4B, cyan). This observation suggests that SST-INs are likely recruited by thalamic inputs for feedforward inhibition of PT neurons via α5-GABARs^44^, which modulate thalamic synaptic excitation at subthreshold membrane potentials. Subsequent blockade of NMDARs with D-APV (50 μM) reduced the peak amplitude and the integral of PSPs to baseline levels (Figure 4B, red versus black traces). This observation indicates that, in the healthy state, the subthreshold PSP was mediated primarily by non-NMDARs, with minimal involvement of NMDARs (Figure 4C). Since NMDAR blockade abolished the effects of α5IA, it suggests that α5-GABARs preferentially modulate NMDAR function and have little impact on non-NMDARs in the healthy state.

**Figure 4.**
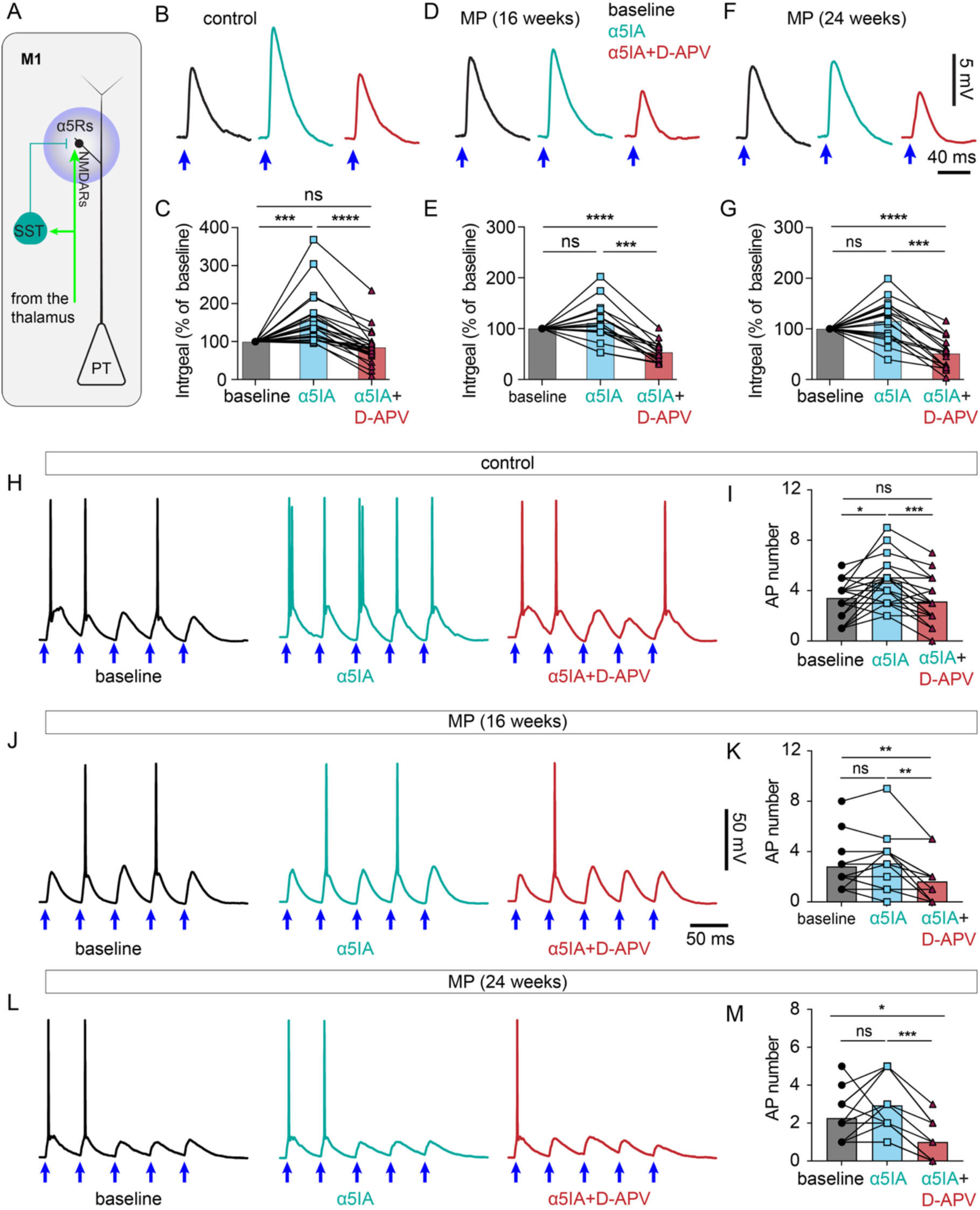
Weakened α5-GABA_A_R modulation of NMDAR activation at the thalamo-PT synapses during progressive loss of SNc DA neurons. **A)** Experimental design. Optogenetics was used to selectively activate thalamic inputs to PT and SST-INs in M1 in order to study the interaction between α5-GABARs and NMDARs in PT neurons. **B-C**) Effects of α5IA (cyan) and D-APV (red) on postsynaptic potentials in PT neurons from control mice. Postsynaptic potentials were recorded sequentially at baseline, in the presence of α5IA (% baseline, 155.8±13.5%, p = 0.0009 versus baseline), and α5IA plus D-APV (% baseline, 84.7±9.14%, p < 0.0001 versus α5IA, Friedman test followed by Dunn’s tests, n = 24 cells/6 mice). **D-G**) Effects of α5IA (cyan) and D-APV (red) on postsynaptic potentials in PT neurons from MP mice at 16 weeks of age (D-E) and 24 weeks of age (F-G). At 16 weeks of age, α5IA = 111.2±9.9%, p > 0.99 versus baseline; α5IA+D-APV = 54±4.9%, p < 0.0001 versus α5IA, n = 16 cells/3 mice. At 24 weeks of age, α5IA = 115.4±9.9%, p > 0.99 versus baseline; α5IA+D-APV = 51.4±7.8%, n = 17 cells/4 mice, p < 0.0001 versus α5IA. Friedman test followed by Dunn’s tests. **H-I**) Effects of α5IA (cyan) and D-APV (red) on synaptically-driven AP firing in PT neurons from control mice. Optogenetic stimulations (blue arrows) were delivered at 20 Hz to evoke action potentials and postsynaptic potentials at resting membrane potential. AP numbers, baseline = 3.4±0.3, α5IA = 4.6±0.3, α5IA+D-APV = 3.1±0.3, n= 27 neurons/6 mice, Friedman test followed by Dunn’s tests. **J-M**) Effects of α5IA (cyan) and D-APV (red) on synaptically-driven AP firing in PT neurons from MP mice at 16 weeks of age (J, K) and 24 weeks of age (L-M). At 16 weeks, baseline = 2.8±0.5, α5IA = 3.1±0.5, α5IA+D-APV = 1.6±0.4, n = 16 neurons/3 mice. At 24 weeks, baseline = 2.3±0.3, α5IA = 2.9±0.4, α5IA+D-APV = 1.0±0.3, n= 15 neurons/4 mice. Friedman test followed by Dunn’s tests. ns, not significant, *p < 0.05, **p < 0.01, ***p < 0.001, ****p < 0.0001.

In 16- and 24-week-old MP mice, we found that α5IA failed to elicit a robust effect on the optogenetically evoked subthreshold PSPs of PT neurons (Figure 4D-G, cyan versus black traces in D and F). This indicates that α5-GABARs in PT neurons were barely recruited by thalamic inputs in MP mice, in striking contrast to their pronounced activation in the healthy state (Figure 4B). Subsequent NMDAR antagonism by D-APV (50 μM) significantly decreased the peak amplitude and integral of the subthreshold PSPs to a level below the baseline in MP mice of both 16- and 24-weeks of age (Figure 4D-G, black and red traces in D and F). This observation indicates that in parkinsonian states, NMDARs were persistently activated at thalamo-PT synapses even at baseline, concurrent with impaired α5-GABAR-mediated inhibition (Figure 3).

To further understand the functional impact of impaired α5-GABARs on suprathreshold response of PT neurons (i.e., neuronal firing), we delivered a train of optogenetic stimulation at 20 Hz to evoke synaptically-driven APs (Figure 4H). In healthy control mice, thalamic input could effectively evoke AP firing in M1 PT neurons, consistent with an early report that synchronous thalamic inputs provide potent excitation that drives cortical firing^19,28^. Inhibition of α5-GABARs significantly boosted the effectiveness of thalamus-driven AP firing of PT neurons, as reflected by the increased AP numbers after α5IA application (Figure 4H, I, black versus cyan traces). In the presence of D-APV (50 μM), the effect of α5IA in facilitating AP firing was abolished (Figure 4H, I, red and black traces). These data suggest that α5-GABAR antagonism facilitates AP firing mainly by enhancing NMDAR function. However, in 16- and 24-week-old MP mice, α5IA failed to increase AP firing in PT neurons (Figure 4J-M, black versus cyan traces). Subsequent blockade of NMDARs by D-APV reduced AP number to a level below baseline (Figure 4J-M, red and black traces). These data suggest that impaired α5-GABAR-mediated inhibitory control of PT neurons is associated with overactivation of NMDARs at thalamo-PT synapses.

Furthermore, we injected the AAV-ChR2(H134R)-eYFP into the sensory cortex to study the impact of α5-GABARs on corticocortical inputs to M1 PT neurons (i.e., inputs from the sensory cortex). Strikingly, application of α5IA or D-APV did not alter the integral of PSPs (i.e., the area under the curve) evoked by optogenetic stimulation of cortical afferents to M1 (Figure S6). These observations indicate that sensory cortical inputs are less effective in activating M1 SST-INs and the associated α5-GABAR signaling and that NMDARs do not predominantly contribute to the subthreshold synaptic responses at the sensory cortical input to PT neurons. Compared with thalamic inputs (Figure 4), these input-specific responses might reflect a biased recruitment of M1 inhibitory microcircuits by the sensory cortical inputs^44^ and/or by the compartment-specific levels of NMDAR expression in cortical pyramidal neurons^45^. Together, these data suggest that during chronic loss of SN DA neurons, impaired α5-GABAR-mediated inhibition promotes NMDAR activation and facilitates the generation of thalamus-driven APs in PT neurons.

Taken together, these data suggest that SST-INs tightly control NMDAR-mediated thalamic excitation of PT neurons via the slow-kinetic α5-GABA_A_Rs under healthy conditions. During progressive loss of SNc DA neurons, diminished α5-GABAR-mediated inhibition leads to persistent NMDAR overactivation at thalamo-PT synapses, which likely plays a critical role in triggering synaptic and cellular deficits in PT neurons.

### GluN2B-NMDARs contribute to the reduced thalamo-PT synaptic dysfunction

Synaptic GluN2B subunits are highly expressed in prefrontal cortical pyramidal neurons in both rodents and nonhuman primates ^46,47^, critically contributing to sustained neuronal firing during working memory tasks. The results described above suggest that the slow-kinetic GluN2B-NMDARs in PT neurons become overactivated, perhaps contributing to the overall weakened AMPAR-mediated thalamo-PT synaptic transmission after the loss of SNc DA neurons. To test this hypothesis, we studied the effects of knocking down GluN2B expression in PT neurons from both control and MP mice at 24 weeks of age to determine whether this manipulation can ameliorate thalamo-PT synaptic deficits. Selective GluN2B knockdown *in vivo* was achieved using AAV-based CRISPR/Cas9 technologies^48^. We injected the retrogradely transported AAVrg-flpo into the pons and AAV-FLEXfrt-SaCas9-U6-sgGrin2b or AAV-FLEXfrt-SaCas9-U6-sgRosa26 as controls into M1 (Figure 5A). AAV-FLEXfrt-SaCas9-U6-sgGrin2B or AAV-FLEXfrt-SaCas9-U6-sgRosa26 was mixed and co-injected with AAV-fDIO-tdTomato for visualization of injection sites and subsequent functional validation^48^.

**Figure 5.**
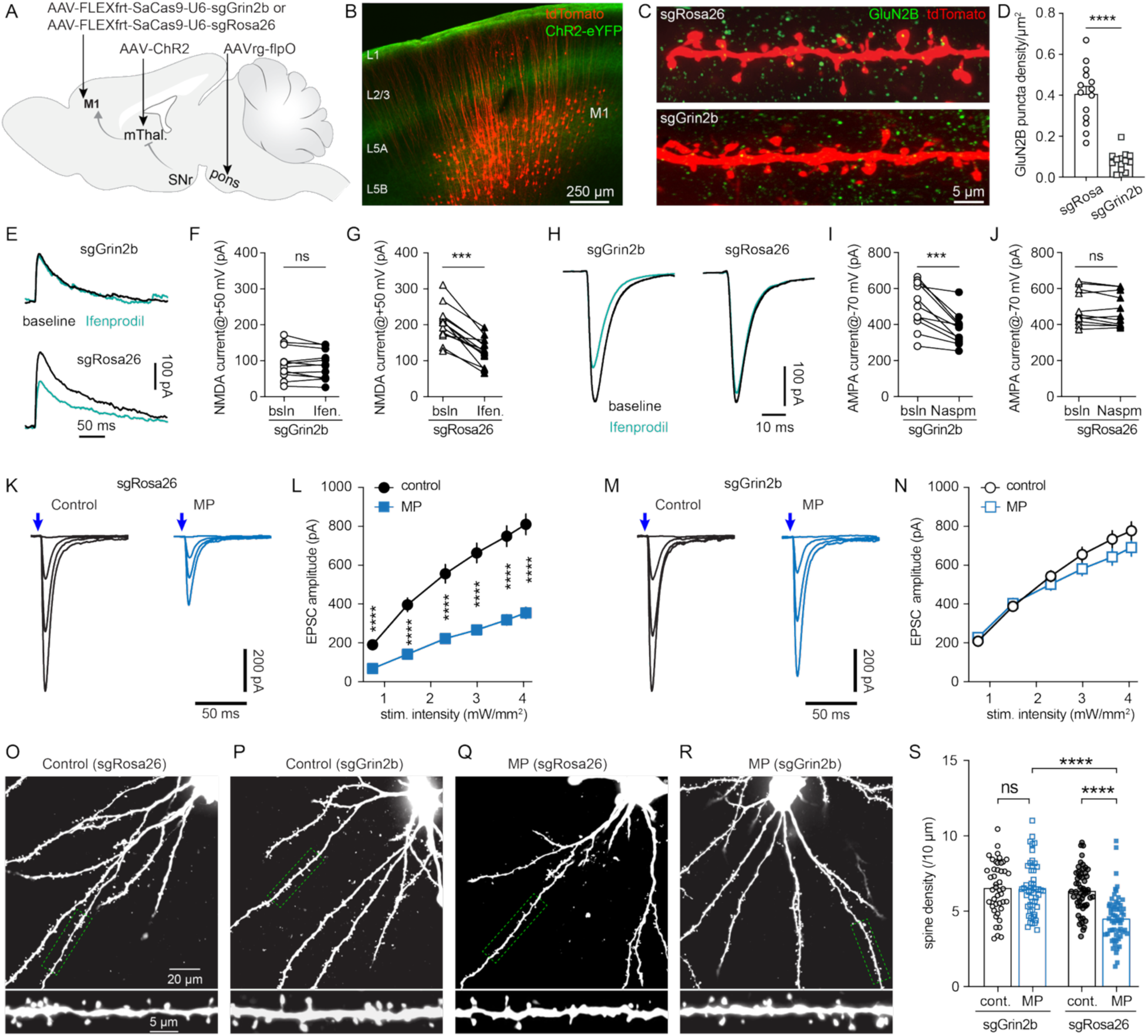
GluN2B knockdown rescues the impaired thalamo-PT synaptic connections in advanced MP mice. **A**) Experimental design. **B**) Visualization of retrogradely labeled PT neurons in layer 5 of M1 by tdTomato. **C-D**) CRISPR-mediated GluN2B knockdown from PT neurons. GluN2B puncta, sgRosa26 = 0.48±0.04 puncta/μm^2^, n = 24 segments/4 mice, sgGrin2b = 0.10±0.02 puncta/μm^2^, n = 18 segments/3 mice, p < 0.0001, MWU test. **E-G**) GluN2B-selective antagonist ifenprodil (3 μM) decreased the amplitude of NMDAR-mediated EPSPs at +40 mV in PT neurons from mice with sgRosa26 injections (baseline = 230.8±23.3 pA, ifenprodil = 149.9±17.0 pA, n = 16 neurons/7 mice, p < 0.0001, WSR test) relative to those in sgGrin2b (baseline = 92.5±12.7 pA, ifenprodil = 88.3±11.4 pA, n = 12 neurons/3 mice, p = 0.34, WSR test). **H-J**) Effects of GluA2-lacking AMPAR antagonism by NASPM on AMPAR-mediated EPSCs in PT neurons infected with sgGrin2b (baseline = 506.1±38.8 pA, NASPM = 376.3±27.1 pA, n = 11 neurons, p = 0.001, WSR test) and sgRosa26 controls (baseline = 464.6±29.3 pA, NASPM =454.8±28.4 pA, n = 14 neurons/4 mice, p = 0.17, WSR test). **K-L**) The amplitude of thalamo-PT EPSCs in 24-week-old controls and MP mice receiving Rosa26 injections. Control = 48 neurons/5 mice, MP = 61 neurons/6 mice, two-way RM ANOVA followed by Sidak tests. **M-N**) GluN2B knockdown restored the thalamo-PT synaptic strength in MP mice to littermate controls. Control = 40 neurons/4 mice, MP = 40 neurons/4 mice, two-way RM ANOVA followed by Sidak tests. **O-S**) GluN2B knockdown rescues the dendritic spine loss in PT neurons of MP mice relative to controls. Control/sgRosa26 =6.34±0.16 spine/10 μm, n = 82 segments/6 mice; MP/sgRosa26 = 4.43±0.17 spine/10 μm, n =81 segments/6 mice; p < 0.0001; control/sgGrin2b = 6.57±0.26 spines/10 μm, n = 44 segments/4 mice, MP/sgGrin2b = 6.51±0.25 spines/10 μm, n = 49 segments/4 mice, p > 0.99. Two-way ANOVA followed by Sidak’s test.

The effectiveness of GluN2B knockdown was confirmed using both immunohistochemistry and *ex vivo* electrophysiology. Immunohistology showed a robust reduction of GluN2B immunoreactivity in PT neurons from animals receiving AAV-FLEXfrt-SaCas9-U6-sgGrin2b injection relative to AAV-FLEXfrt-SaCas9-U6-sgRosa26 or AAV-fDIO-tdTomato-injected controls (Figure 5B-D). In *ex vivo* brain slices, while the selective GluN2B-NMDARs antagonist Ifenprodil (3 μM)^49^, could significantly and consistently reduce the amplitude of NMDAR-mediated oEPSC in PT neurons from AAV-FLEXfrt-SaCas9-U6-sgRosa26-injected controls (Figure 5E-G), this effect was absent in PT neurons from AAV-FLEXfrt-SaCas9-U6-sgGrin2b-injected mice (Figure 5E-G). The above results confirm the effective and robust reduction of GluN2B-containing NMDARs in PT neurons infected with sgGrin2b. In addition, we found that AMPAR-mediated oEPSCs at the thalamo-PT synapse exhibit strong inward rectification (data not shown), indicating alterations in postsynaptic AMPAR composition. Consistently, 1-naphthylacetyl spermine (NASPM, 100 μM)^35,50^, a selective antagonist of GluA2-deficient, Ca^2+^- permeable AMPARs, significantly decreased the amplitude of AMPA-oEPSCs in PT neurons with GluN2B knockdown (Figure 5H-I). In contrast, NASPM did not alter the amplitude of AMPAR-mediated oEPSCs in PT neurons with sgRosa26 expression (Figure 5H, J). These results suggest that GluN2B subunit knockdown increases the function of GluA2-deficient, Ca^2+^-permeable AMPARs in PT neurons.

To investigate the effect of GluN2B knockdown on thalamo-PT synaptic strength, we measured the amplitude of AMPAR-mediated oEPSCs at -80 mV from the thalamo-PT synapses in MP mice and littermate controls receiving AAV9-FLEXfrt-SaCas9-U6-sgRosa26 or AAV9-FLEXfrt-SaCas9-U6-sgGrin2b injections into M1, together with AAVrg-flpo injection into the pons. In SaCas9-U6-sgRosa26-injected mice, there was a significant reduction in the amplitude of oEPSCs from 24-week-old MP mice relative to littermate controls (Figure 5K-L). However, in GluN2B knockdown mice, we found no difference in the amplitude of oEPSCs between MP and control mice (Figure 5M-N).

Morphologically, there was a robust loss of dendritic spines of PT neurons in MP mice relative to littermate controls (Figure 5O-S), consistent with the observation from parkinsonian mice and monkeys^19,42^. Notably, while GluN2B subunit knockdown produced a negligible effect on dendritic spine density of PT neurons in controls, it restored the spine density of PT neurons in MP mice relative to Rosa26-injected controls (Figure 5S). Together, these results suggest that suppression of the overactivated GluN2B-containing NMDARs can ameliorate thalamo-PT synaptic deficits after loss of SNc DA neurons.

## L-DOPA ameliorates thalamocortical synaptic deficits in MP mice

In parkinsonian states, the hypersynchronous rhythmic GABAergic output from the basal ganglia abnormally entrains thalamic neurons, resulting in pathologically synchronized burst pattern of firing^16,17,51^. Synchronous glutamatergic synaptic inputs from the thalamus can effectively drive cortical neuronal firing^28^. Thus, it is reasonable that after DA loss, persistently synchronous thalamocortical inputs drive overstimulation of dendritic GluN2B-NMDARs in PT neurons, triggering maladaptive downregulation of thalamo-PT synapses, as in the subthalamic nucleus^35^. In this case, the impairment of neurotransmission at thalamo-PT synapses may be preventable or rescuable by suppressing pathological basal ganglia output. We sought to test this hypothesis.

Early work showed that the beneficial effect of L-DOPA may involve the desynchronization of basal ganglia output by reducing neuronal firing rate^52^, disrupting abnormally synchronized burst firing^53^, or suppressing oscillatory activity^54–56^ in PD patients and parkinsonian animals. Thus, we first examined whether L-DOPA treatment can concurrently restore the thalamocortical connections under parkinsonian conditions. MP mice and littermate controls at 21 weeks of age received Retrobeads injections into the pons to retrogradely label PT neurons and AAV9-hSyn-ChR2(H134R)-eYFP injection into the motor thalamus to selectively stimulate thalamocortical projections (Figure 6A). Once the animals had recovered from surgery, they received L-DOPA or saline injections (4 mg.kg, i.p., twice/day) followed by daily locomotion tests for 5 consecutive days (Figure 6B). *Ex vivo* brain slice recordings were conducted ∼ 3 hours after the last L-DOPA injection. As expected, saline-injected MP (MP/saline) mice showed significantly impaired locomotor activity in an open field arena relative to saline-injected littermates (littermate/saline) across days (Figure 6C, E), reflecting their parkinsonian motor deficits. Interestingly, L-DOPA-injected MP (MP/L-DOPA) mice showed gradually improving locomotor activity over the course of treatment, reaching levels comparable to L-DOPA-injected controls (littermate/L-DOPA) on days 4 and 5 (Figure 6D, F). Specifically, MP/L-DOPA mice showed similar distance traveled relative to littermate/L-DOPA mice from day 3 to day 5 (Figure 6F). MP/L-DOPA mice did not manifest dyskinesia-like behavior (supplementary video 1)^57,58^. These observations indicate that plasticity processes were induced by chronic L-DOPA treatment and are consistent with the prior reports on the responsiveness of MP mice to L-DOPA injections^23,25^.

**Figure 6.**
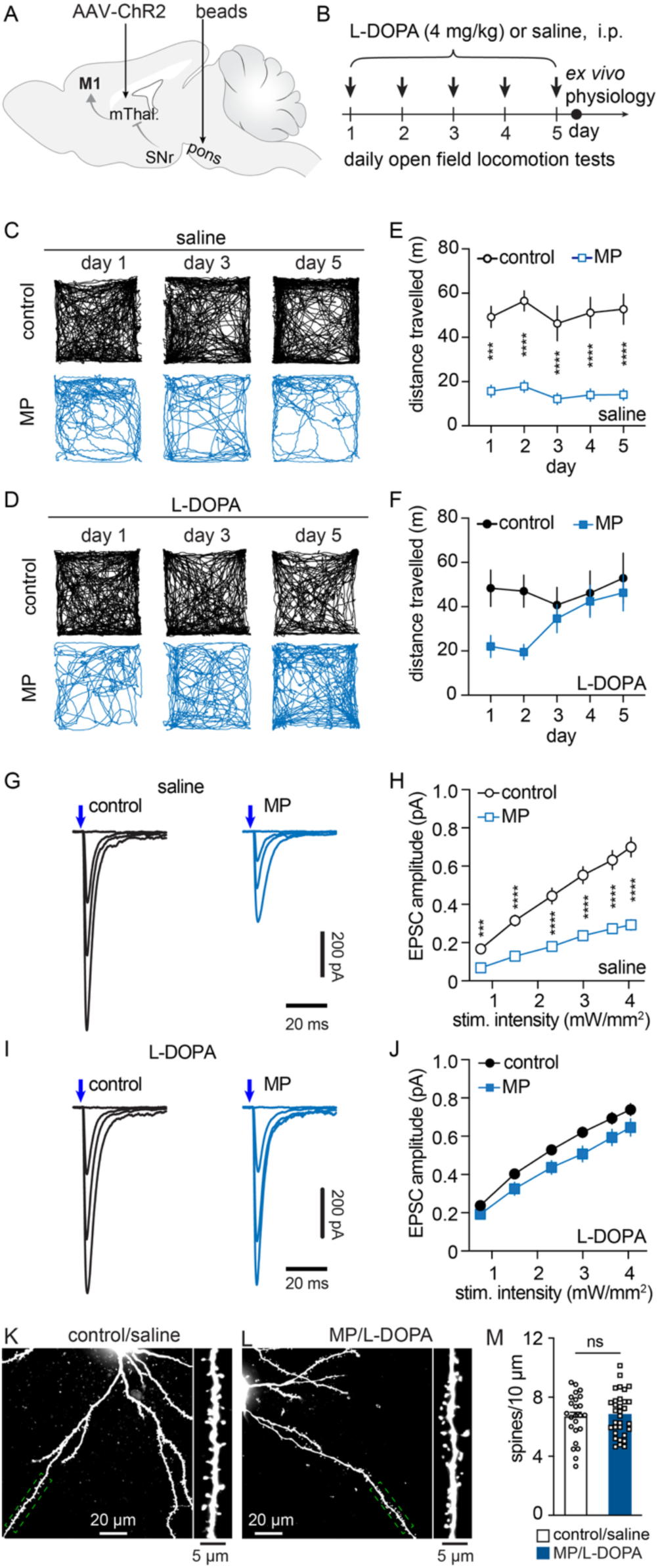
Chronic L-DOPA treatment ameliorates thalamocortical synaptic deficits in advanced MP mice. **A**) Experimental design. **B**) L-DOPA treatment paradigm as well as experimental design of behavioral and physiological studies. *Ex vivo* electrophysiology studies were conducted after the last L-DOPA injection and open field locomotion test on day 5. **C**) Track plots showing locomotor activities of control (top panel) and MP mice (bottom panel) that were injected with saline (C) or L-DOPA (D) across days. **E-F**) Summary results of locomotor activities of saline- (E) and L-DOPA- (F) injected control and MP mice across days. control/saline = 10 mice, MP/saline = 8 mice; control/L-DOPA = 10 mice, MP/L-DOPA = 11 mice. Two-Way RM ANOVA followed by Sidak tests. **G-H**) Saline-injected MP mice showed reduced amplitude of thalamo-PT EPSCs relative to saline-injected controls. Control = 22 neurons/4 mice, MP = 21 neurons/4 mice. Two-Way RM ANOVA followed by Sidak tests. **I-J**) Restoration of the amplitude of thalamo-PT EPSCs in L-DOPA-injected MP mice relative to L-DOPA-injected controls. Control = 29 neurons/5 mice, MP = 18 neurons/3 mice. Two-Way RM ANOVA followed by Sidak tests. **K-M**) Chronic L-DOPA treatment restores spine loss of PT neurons in advanced MP mice. spine density: littermate/saline = 6.66±0.32 spines/10 μm, n = 24 segments/3 mice; MP/L-DOPA = 6.83±0.26 spines/10 μm, n = 32 segments/3 mice, p = 0.9, MWU. ns, not significant, *p < 0.05, ***p < 0.001, ****p < 0.0001.

Next, we performed *ex vivo* brain slice recordings to determine whether chronic L-DOPA treatment could ameliorate the impaired thalamo-PT synaptic connections. As expected, we found that MP/saline mice showed significantly reduced amplitude of thalamo-PT oEPSCs relative to littermate/saline mice at 24 weeks of age (Figure 6G, H). After chronic L-DOPA treatment, MP/L-DOPA mice showed a comparable amplitude of thalamo-PT oEPSCs relative to littermate/L-DOPA mice (Figure 6I, J).

Loss of midbrain DA neurons induces a significant loss of dendritic spines of M1 PT neurons in both 6-OHDA-lesioned mice^19^ and advanced MP mice (Figure 5O, Q, S). Thus, we determined if chronic L-DOPA treatment could rescue dendritic spine loss in MP mice. We quantified the dendritic spines of biocytin-labeled PT neurons and found no difference in dendritic spine density in PT neurons between MP/L-DOPA and littermate/saline mice (Figure 6K, M). These data suggest that chronic L-DOPA treatment also rescues dendritic spine loss. Combining electrophysiological and morphological results, we demonstrated that chronic L-DOPA treatment ameliorates the impaired thalamic inputs to PT neurons in parkinsonism.

To exclude the possibility that L-DOPA’s effect on thalamocortical synapses is mediated by direct modulation of dopaminergic receptors in M1, we adopted a chemogenetic approach to directly suppress basal ganglia output (without pharmacological stimulation of cortical DA receptors) and examined its effect on thalamo-PT synapses in late-stage MP mice and littermate controls. We injected AAVrg-flpo into the motor thalamus to retrogradely express FlpO in the thalamus-projecting SNr neurons and AAV9-hSyn-fDIO-hM4Di(Gi)-mCherry into the SNr of MP mice and littermates at 21 weeks of age (Figure S7A-B). Application of a potent and selective DREADDS agonist, deschloroclozapine (DCZ, 1 μM), effectively suppressed the autonomous firing of hM4Di-expressing SNr cells (Figure S7C-D). Next, we employed a 5-day chemogenetic treatment paradigm, as in chronic L-DOPA studies (Figure 6B), to chronically suppress SNr neuronal activity and then examined its impact on animal behavior. We found that both saline-injected littermate controls (littermate/saline) and saline-injected MP mice (MP/saline) showed reduced locomotor activity post-injection compared with pre-injection levels, likely due to habituation to the testing environment (Figure S7E, G). There was no group difference in the magnitude of movement reduction across days (Figure S7G). In contrast, while DCZ-injected littermate controls (littermate/DCZ) exhibited a generally reduced post-injection locomotor activity, the overall locomotor movement of DCZ-injected MP mice (MP/DCZ) was significantly increased relative to pre-injection baseline (Figure S7F, G). These observations support the conclusion that inhibitory DREADDs can effectively suppress basal ganglia output *in vivo*, thereby improving motor function in parkinsonian mice.

We next assessed whether chronic chemogenetic suppression of SNr activity ameliorates the impaired thalamo-PT synaptic strength in MP mice. After 5 days of injections, MP/saline mice exhibited reduced amplitude AMPAR-mediated oEPSCs at thalamo-PT synapses relative to littermate/saline mice (Figure S7H, I), consistent with the impaired thalamic projection to M1 PT neurons after the degeneration of nigrostriatal DA projections. In contrast, the amplitude of AMPAR-mediated oEPSCs at thalamo-PT synapses was comparable between MP/DCZ mice and littermate/DCZ mice (Figure S7J, K). Given that cortical DA receptors were not stimulated in the above chemogenetic studies, we conclude that suppression of BG output is sufficient to rescue thalamo-PT synaptic strength under parkinsonian conditions.

## Early L-DOPA treatment prevents thalamocortical synaptic deficits during progressive parkinsonism

The experiments described above demonstrated that M1 circuit function remains intact until a moderate stage of Parkinsonism (e.g., 16-week-old MP mice), which is associated with >80% striatal DA depletion (Figures 1 and S1). A corollary prediction is that cortical circuit impairments in parkinsonism could be halted once the development of abnormal basal ganglia firing is suppressed, for example, by early use of dopaminergic medications. To determine the effects of early L-DOPA administration on the functions of cortical circuits, we retrogradely labeled PT neurons in M1 and expressed ChR2 in the thalamus of 8-week-old MP mice and littermate controls, using the methods mentioned above (Figure 7A-B). Three to 4 weeks later (i.e., at 12 weeks of age), MP mice and littermate controls were treated with L-DOPA (4 mg/kg, twice/day for 4 weeks, i.p.), which did not induce dyskinesia-like behavior. We then examined thalamo-PT synaptic strength at 16 weeks of age (Figure 7B). At 16 weeks, we found that daily L-DOPA treatment did not affect the extent of striatal DA depletion in MP mice (Figure 7C, D). This observation indicates that early L-DOPA administration does not alter the temporal progression of dopaminergic neurodegeneration, as suggested by the literature on PD patients^59–61^.

**Figure 7.**
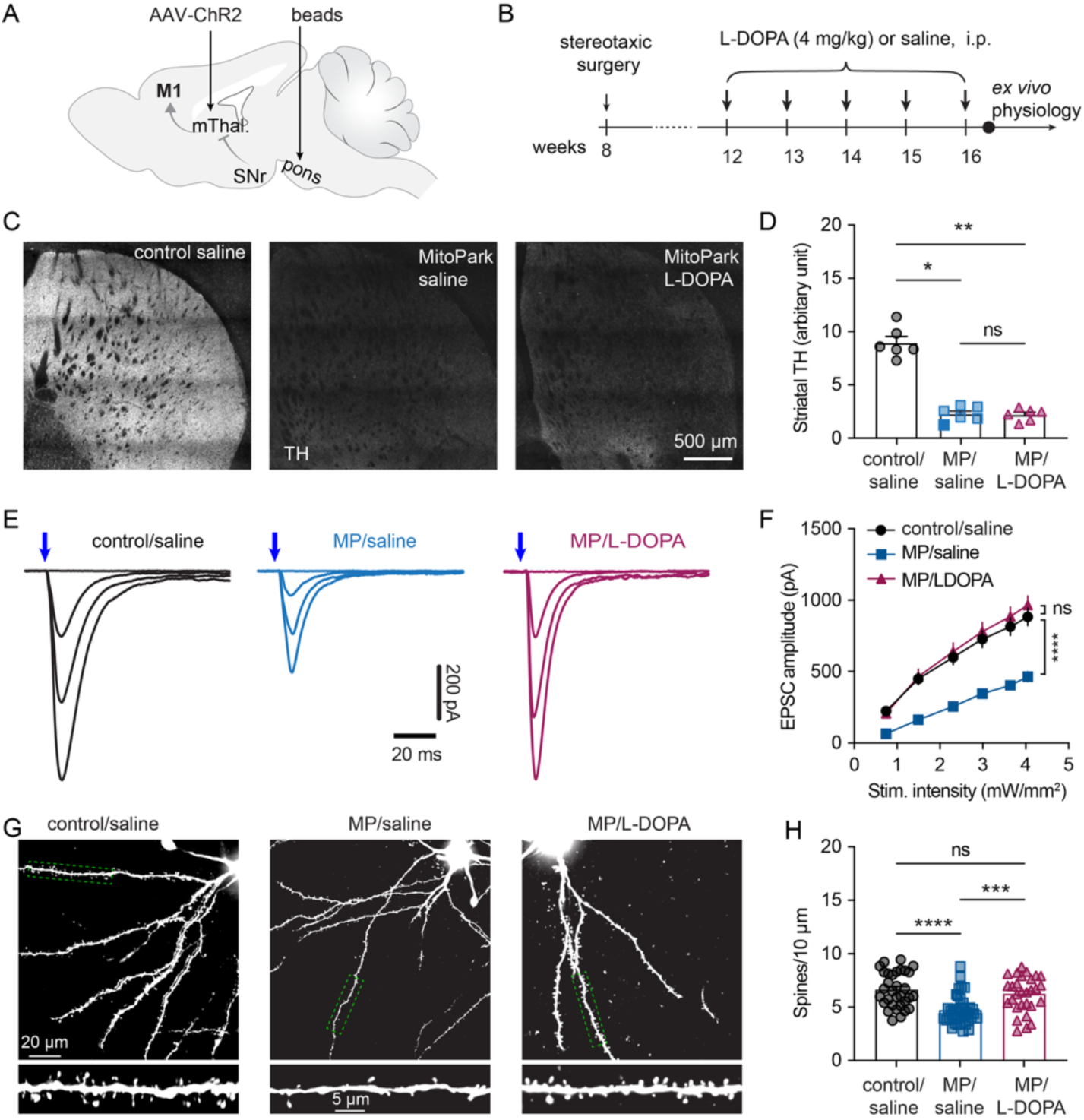
**Early L-DOPA treatment can delay thalamo-PT synaptic deficits during the progressive degeneration of SNc DANs. A-B**) Experimental design. **C-D**) Confocal images (C) and summary graph (D) showing TH-ir in the dorsal striatum of saline-injected controls (8.97±0.59, n= 6 slices/3 mice), saline-injected MP mice (2.28±0.29, n= 6 slices/3 mice), and L-DOPA-injected MP mice (2.21±0.24, n= 6 slices/3 mice) at 16 weeks of age. **E-F**) Representative traces of the thalamo-PT oEPSCs and summarized results of thalamo-PT EPSC amplitude of control/saline (28 neurons/3 mice), MP/saline(30 neurons/3 mice), and MP/L-DOPA (28 neurons/3 mice) mice at 16 weeks of age. Two-way RM ANOVA followed by Sidak test. *Post hoc* analysis results are available in the Source Data Table. **G-H**) Representative confocal images of dendritic spines under low and high magnification (G) and summarized results (H) of dendritic spine density in control/saline (6.6±0.29/10 μm, 32 segments/3 mice), MP/saline mice (4.7±0.21/10 μm, 39 segments/3 mice), and MP/L-DOPA mice (6.2 ± 0.31/10 μm, 29 segments/3 mice) at 16 weeks of age. ***p < 0.001, ****p < 0.0001, Kruskal-Wallis test.

To determine the impact of 4 weeks of L-DOPA treatment on M1 circuit adaptation at 16 weeks of age, the thalamo-PT synaptic connection strength was examined in control and MP mice that received saline injections (MP/saline), as well as MP mice that received L-DOPA injections (MP/L-DOPA). A profound reduction in the amplitude of thalamo-PT oEPSCs was found in MP/saline mice relative to littermate/saline mice (Figure 7E, F), which is consistent with the earlier observations. In MP/L-DOPA mice, the amplitude of thalamo-PT EPSCs was increased significantly and normalized to the level of littermate/saline mice (Figure 7E, F). We also found that MP/saline mice exhibited a significant reduction in the dendritic spine density relative to littermate/saline mice (Figure 7G, H). However, such differences were absent in MP/L-DOPA mice relative to control/saline group (Figure 7G, H). These results suggest that early L-DOPA administration delays certain aspects of cortical circuit adaptations during Parkinsonism, although it has a negligible effect on the trajectory of dopaminergic neurodegeneration.

## Discussion

In the present work, we employed a genetic model of gradual dopaminergic neurodegeneration and defined the time course of synaptic and cellular plasticity of M1 circuits during progressive parkinsonism. We demonstrated that SST-INs exert asymmetric innervation of PT and IT neurons in M1 via α5-GABAR-mediated inhibition under physiological conditions (Figure 8A). Using both neurotoxin-based and genetic parkinsonian models, we demonstrated that loss of SNc DA neurons reduces α5-GABAR-mediated inhibition of PT neurons. This attenuation impairs gating of dendritic NMDAR function by α5-GABARs, which at least partially contributes to cell-type-specific cortical alterations in parkinsonism (Figure 8B). Moreover, the altered thalamic inputs to M1 are associated with a substantial loss of dendritic spines in M1 PT neurons. We also demonstrated that the functional and morphological alterations of thalamic inputs to M1 PT neurons can be ameliorated by L-DOPA treatment, perhaps by suppressing abnormal basal ganglia outputs in parkinsonism.

**Figure 8.**
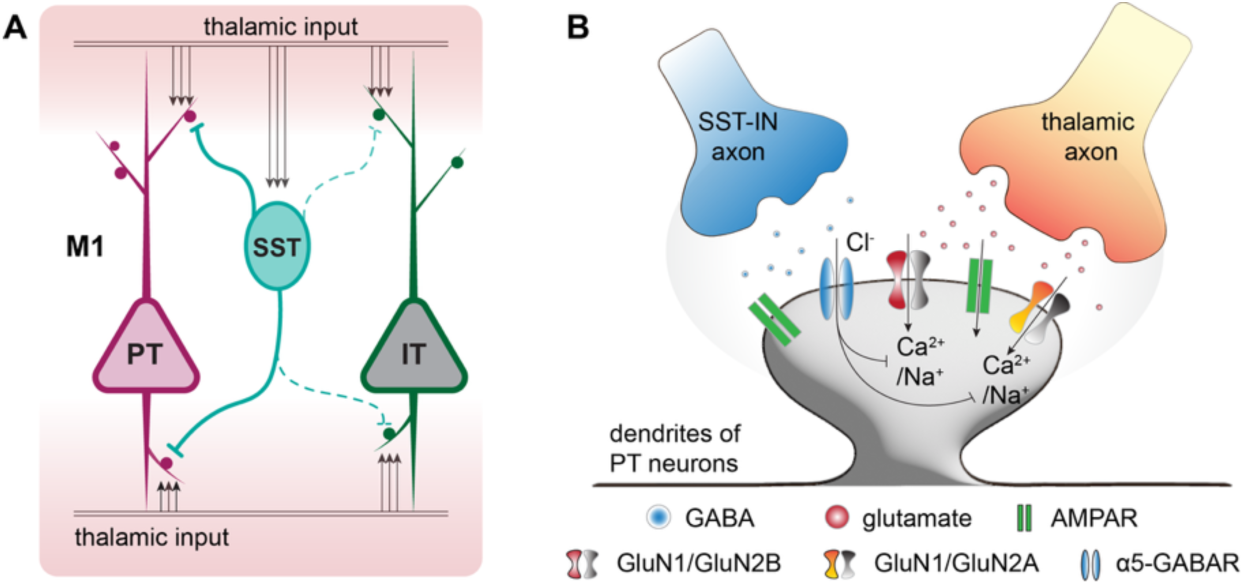
Graphic summary. **A**) SST-INs exert asymmetric inhibition of PT (solid lines) and IT (dashed lines) neurons in physiological states. **B**) Cartoons showing key microcircuits underlying M1 dysfunction, involving the interaction between α5-GABARs and NMDARs at dendritic spines of M1 PT neurons.

## Cellular and synaptic adaptation in parkinsonian conditions

Evidence from rodent and NHP studies suggests that loss of dopaminergic neuromodulation triggers complex circuit adaptations in the cortico-basal ganglia-thalamocortical network in PD, which partially contribute to the development of motor and nonmotor symptoms in patients^1,5,62^. However, compared with the well-studied basal ganglia nuclei, we have far less knowledge of cerebral cortical dysfunction during progressive parkinsonism, which has long been thought to underlie motor and cognitive deficits in PD ^6,7,63^.

Earlier work on MPTP-treated NHPs reported abnormal anatomical and physiological properties of M1 circuits following nearly complete degeneration of SNc DA neurons^12–14,42^. Of particular interest, Pasquereau et. al. reported that cortical PT neurons were selectively affected in MPTP-treated parkinsonian monkeys, whereas the activity of corticostriatal neurons (corresponding to IT neurons in rodents) was largely unaltered^13^. Such cell-subtype-selective alterations suggest specific neural plasticity processes are triggered within cortical circuits. Supporting this concept, Villalba et. al. reported cortical layer-specific anatomical changes of thalamocortical synapses in M1 of MPTP-treated monkeys, including a robust reduction of vGluT2-ir (a marker of thalamic axon terminals) in the deeper layers, but not the superficial layer of M1^42^. Recently, we studied the cellular and synaptic changes in M1 circuits after severe degeneration of midbrain DA neurons in mice with 6-OHDA lesions. Our work suggested that the intrinsic excitability and thalamic excitation of M1 PT neurons are selectively impaired following the SNc DA neurodegeneration^19,20^.

However, the previously used non-human primate and rodent models do not recapitulate the progressive nature of PD. In the present study, we therefore sought to determine the time course of M1 circuit changes and their relationship to the emergence of motor deficits using a progressive mouse model of nigrostriatal neurodegeneration. We found that the physiological adaptations in M1 become detectable after a significant striatal DA depletion (i.e., > 80%), which is associated with a moderate deficit in general motor function (Figures 1 and S1). Gradual DA depletion decreases the thalamic excitation of M1 PT neurons before downregulating their intrinsic excitability (Figure 1). Moreover, these changes are specific to PT neurons: neither thalamic excitation nor intrinsic excitability of IT neurons in M1 is altered at the advanced stage of MP mice (Figure S3), consistent with prior work^13,19^.

## Microcircuit mechanisms of cortical dysfunction

Thalamic axon terminals form synapses with cortical pyramidal neurons primarily on dendritic spines^18,42^, which are often co-innervated by GABAergic synapses^43^. This observation indicates that thalamocortical excitation closely interacts with GABAergic inhibition within a single dendritic spine. Moreover, weakened GABAergic inhibition of the motor cortex has been implicated in PD patients^64–66^ and parkinsonian animals^67,68^, which might contribute to the manifestation of motor deficits, like bradykinesia ^65,67^. We found that PT neurons receive more potent GABAergic inhibition than IT neurons in the healthy state (Figure S5) and that GABAergic inhibition of PT neurons, but not that of IT neurons, decreases following the loss of SNc DA neurons (Figure S5). Furthermore, SST-INs exert potent α5-GABA_A_R-mediated inhibition on M1 pyramidal neurons (Figure 2)^37^. Thus, the recruitment of α5-GABARs by SST-INs to inhibit pyramidal neurons is a conserved feature of cortical and hippocampal circuits^38^. Importantly, the SST inhibition of PT neurons is reduced in both models of parkinsonism (Figure 3). Such a reduction is associated with a decreased presynaptic GABA release probability at SST-PT synapses following loss of SNc DA neurons, which at least in part contributes to the reduced frequency of sIPSCs in PT neurons from both parkinsonian models (Figure 3).

We found that α5-GABAR-mediated inhibition of NMDAR activity at the thalamo- PT synapses was impaired in parkinsonian states, a change detectable at a moderate motor stage (i.e., at 16 weeks of age, Figure 4). This observation provides important insights into our understanding of the cortical pathophysiology of PD. In parkinsonism, synchronous inputs from the basal ganglia-recipient motor thalamus could trigger simultaneous glutamate release^16^, facilitating postsynaptic NMDAR activation. Combined with the impaired α5-GABAR-mediate inhibition, these pathological thalamic inputs likely drive overactivation of postsynaptic NMDARs, particularly the GluN2B-containing NMDARs at the thalamo-PT synapses. We propose that such chronic overstimulation of NMDARs mediates the functional and morphological alterations of thalamocortical synapses through Ca^2+^ signaling-mediated or non-ionotropic mechanisms, as reported in the subthalamus^35,69,70^. Furthermore, using CRISPR-mediated genetic knockdown of the GluN2B subunit of NMDARs, we found that the impaired thalamo-PT synaptic strength can be restored in MP mice with advanced parkinsonism (Figure 5), supporting the view that changes in GluN2B-NMDAR functions at thalamic synapses on PT neurons have a critical role in the development of cortical circuit dysfunction in parkinsonian conditions^19^.

However, the initial molecular events that trigger the diminished α5-GABAR function in PT neurons remain to be determined, including a redistribution of synaptic versus extrasynaptic α5 subunits^71,72^. It is noteworthy that GluN2B NMDARs are enriched at the thalamo-PT synapses^19^, and that their activation also promotes the internalization of α5-GABARs^71^. Thus, it is plausible to posit that the enhanced synchronous firing of thalamic neurons following the striatal DA depletion promotes the activation of GluN2B-NMDARs at the thalamo-PT synapses, subsequently initiating the internalization of α5-GABARs in PT neurons^28^. Thus, a vicious circle may form between hyperactivated NMDARs and impaired dendrite-targeting α5-GABARs at SST-PT synapses during the pathological network activity after the loss of DA. This work opens the door for future studies to determine the relationship between changes in PT and SST neuronal firing and NMDAR activation during progressive loss of DA, using *in vivo* electrophysiological and circuit-manipulation tools. Notably, GluN2B subunits are highly expressed in the postsynaptic density of the prefrontal cortex in rodents and nonhuman primates, contributing to the maintenance of sustained network firing^46,47^. It would be interesting to know whether synaptic GluN2B subunits contribute to the emergence and maintenance of abnormally synchronized burst firing in M1 pyramidal neurons under parkinsonian conditions^12^.

## Cortical circuit dysfunction linked to basal ganglia abnormality

The temporal profiles of M1 dysfunction suggest that a significant loss of SNc DA neurons (e.g., in MP mice at 16 weeks of age) is required to trigger the observed cortical synaptic and cellular adaptations (Figure 1). Notably, mesocortical dopaminergic neurons in the ventral tegmental area of MP mice are more resilient to degeneration, as supported by unaltered DA concentrations in the frontal cortex up to 24 weeks of age^25^. These observations suggest that cortical abnormalities in progressive parkinsonism may be primarily associated with abnormal basal ganglia activity. Consistently, using pharmacological and chemogenetic approaches, we demonstrated that suppression or disruption of basal ganglia outputs can indeed rescue the reduction of thalamic inputs to PT neurons in animals with advanced parkinsonism (Figures 6 and S7). Human studies have documented that dopamine replacement therapies can induce short- and long-duration responses (SDR and LDR, respectively) in PD patients with the possible involvement of neural plasticity in the development of LDR^73–75^. Whether the observed changes in the thalamocortical network are part of LDR associated with L-DOPA treatment remains to be determined.

Clinical studies suggest that early L-DOPA treatment does not modify disease progression in PD patients, as measured by symptom severity and neuropathology^59–61^. Consistently, 4 weeks of L-DOPA treatment starting at 12 weeks of MP mice produced little effect on the temporal trajectory of nigrostriatal dopaminergic neurodegeneration (Figure 7). However, early L-DOPA treatment can prevent the emergence of the reduced thalamo-PT synaptic deficits during progressive loss of DA. These data indicate that early therapeutic intervention could be an effective approach to slow the propagation of pathological activity of the basal ganglia to the rest of the brain, thereby preserving cortical network function critical for both proper motor control and cognition. Although we have focused on L-DOPA’s effects on the interaction between basal ganglia and thalamocortical dysfunction, the additional contribution of cortical DA receptor signaling to the observed changes in parkinsonian states remains to be determined^76^. Together, our findings provide a scientific rationale for preserving cortical function through treatment with L-DOPA during the early phases of PD.

## Materials and Methods

### Animals

Mice were housed up to five animals per cage under a 12/12 h light/dark cycle with access to food and water *ad libitum* in accordance with NIH guidelines for the care and use of animals. All experimental design and procedures were reviewed and approved by the Institutional Animal Care and Use Committee (IACUC) at Georgetown University Medical Center (protocol#: 2024-0001). Adult C57BL6 mice (JAX#: 000664; RRID, IMSR_JAX:000664) of both sexes were purchased from the Jackson Laboratory and used for 6-hydroxydopamine (6-OHDA) lesion studies. MP mice (DAT-Cre^+/-^, Tfam^fl/fl^) were obtained by crossing mitochondrial transcription factor A (Tfam) floxed mice (Tfam^fl/fl^; JAX#, 026123; The Jackson Laboratory; RRID:IMSR_JAX:026123) with Dat-Cre mice (JAX#, 006660; The Jackson Laboratory; RRID:IMSR_JAX:006660) to selectively knock down Tfam from the dopaminergic neurons, resulting in progressive nigrostriatal degeneration. Their age-matched littermates without DAT-Cre (i.e., Tfam^fl/fl^, or Tfam^fl/+^) were used as controls. Genotyping service was provided by TransnetYX (Cordova, TN). MP mice were further characterized by behavioral assays and *post hoc* immunohistochemistry (see below) to ensure the development of Parkinsonian neuropathology and motor deficits. To study GABAergic synaptic strength between SST-INs and PT/IT neurons, homozygous SST-Cre knock-in mice (JAX stock#: 013044; RRID:IMSR_JAX:013044) were crossed with Ai32(RCL-ChR2(H134R)/EYFP) mice (JAX stock#: 024109; RRID:IMSR_JAX:024109) to generate SST-ChR2 mice for experiments. Both male and female mice were used and randomly assigned to experimental groups.

### Stereotaxic surgery

Viral vectors, neurotoxin, or retrograde tracers were stereotaxically injected into targeted brain regions under 1-2% isoflurane anesthesia using an automated stereotaxic instrument (Cat. No. 71000-s, RWD). Buprenorphine (3.25 mg/kg, subcutaneous) was administered for analgesia. Specifically, 6-OHDA (1.0 μl at 3-4 μg/μl, HelloBio) or vehicle was stereotaxically injected into the medial forebrain bundle (MFB, coordinates in mm: anterioposterior - 0.7; mediolateral +1.2; dorsoventral -4.9) to induce degeneration of the midbrain dopaminergic neurons or serve as vehicle-injected controls, respectively. Intraperitoneal (IP) injections of desipramine (25 mg/kg) and pargyline (50 mg/kg) were performed ∼40 min before the start of the infusion to enhance the selectivity and toxicity of 6-OHDA, respectively. To study the thalamocortical synaptic transmission and plasticity, AAV9-hSyn-ChR2(H134R)-eYFP (RRID: Addgene_26973, 300 nl at 3.6 × 10^12^ GC/ml) was injected into the motor thalamus (i.e., ventromedial and ventrolateral subregions, coordinates in mm: anterioposterior -1.3; mediolateral +0.8; dorsoventral - 4.3). To label PT and IT neurons in M1, Retrobeads (300 nl at 10× dilution, Lumafluor Inc.) or fastblue (300 nl at 5x dilution, Polyscience, Polyscience_17740-1) were injected into the ipsilateral pontine nuclei (coordinates from the bregma in mm, anterioposterior = −5.0, mediolateral = +0.6, dorsoventral = −5.0) or contralateral striatum (coordinates from the bregma in mm, anterioposterior = +0.7, mediolateral = −2.3, dorsoventral =−2.9).

To specifically knockdown GluN2B subunit of PT neurons, pAAVrg-EF1a-Flpo (1:2 dilution, 2.3 × 10^13^ GC/ml, RRID: Addgene_55637) was injected into the pons (anterioposterior = -5.0, mediolateral = ±0.65, dorsoventral = -5.0 from bregma in mm), the mixture of AAV9-FLEXfrt-SaCas9-U6-sgGrin2b (1:5 dilution, 200 nl, Vector biolabs_25110#53, 5.32 × 10^12^ GC/ml) with AAV-EF1α-fDIO-tdTomato (1:20 dilution, 200 nl, RRID: Addgene_128434) was injected into M1 (AP = +1.0 and +1.6, ML = ±1.5, DV = -0.9 from dura). A mixture of AAV-FLEXfrt-sgRosa26 (1:5 dilution, 0.9 × 10^12^ GC/ml)^48^ with AAV-fDIO-tdTomato (1:20 dilution) was used as the control group. Physiological and anatomical studies were performed 3-4 weeks post-stereotaxic surgeries. Detailed procedures of the stereotaxic injection can be found on Protocol.io (dx.doi.org/10.17504/protocols.io.rm7vzye28lx1/v1).

### Brain slice preparation for electrophysiology

All brain slice physiology studies were conducted 3-4 weeks post-injections. To prepare brain slices, mice were anesthetized with ketamine/xylazine and perfused transcardially using ice-cold sucrose-based artificial cerebrospinal fluid (ACSF), equilibrated with 95% O_2_/5% CO_2_ and containing (in mM): 26 NaHCO_3_, 230 sucrose, 10 glucose, 10 MgSO_4_, 2.5 KCl, 1.25 NaH_2_PO_4_, 0.5 CaCl_2_, 1 Na-pyruvate, and 0.005 L-glutathione. 250 μm coronal brain sections containing the M1 were prepared using a vibratome (VT1200s, Leica, RRID:SCR_018453) in the sucrose-based solution at ∼4°C maintained using a recirculating chiller (FL300, Julabo, Allentown, PA). Slices were then transferred to ACSF equilibrated with 95% O_2_/5% CO_2_ and containing (in mM) 126 NaCl, 26 NaHCO_3_, 10 glucose, 2 MgSO_4_, 2.5 KCl, 1.25 NaH_2_PO_4_, 1 Na- pyruvate, and 0.005 L-glutathione at 35°C for 30 min for recovery and then at room temperature until electrophysiology recordings. Details of slice preparation can be found on Protocols.io (dx.doi.org/10.17504/protocols.io.36wgqj2eovk5/v1).

### *Ex vivo* electrophysiology recordings

Brain slices were transferred to a recording chamber and perfused continuously with a recording solution containing (in mM): 126 NaCl, 26 NaHCO_3_, 10 glucose, 3 KCl, 1.6 CaCl_2_, 1.5 MgSO_4_, and 1.25 NaH2PO_4_. The solution was equilibrated with 95% O_2_/5%CO_2_ and maintained at 32-34°C using a controlled in-line heater (TC-324C, Warner Instruments). In addition, to study synaptic strength, TTX (1 μM)/4-AP (100 μM)/GABAzine (10 μM) were included isolate and measure glutamatergic neurotransmission, or TTX (1 μM)/4-AP (100 μM)/DNQX (20 μM)/D-APV (50 μM) were included to isolate and measure GABAergic neurotransmission from the SST-INs to cortical pyramidal neurons. To assess the intrinsic excitability, DNQX (20 μM)/D-APV (50 μM)/GABAzine (10 μM) were included to block ionotropic glutamatergic and GABAergic receptors. Neurons were visualized using a charge-coupled device camera (SciCam Pro, Scientifica, UK) and a SliceScope Pro 6000 system (Scientifica, UK) controlled by a motorized micromanipulator. Retrogradely labeled pyramidal neurons in M1 were targeted for whole-cell patch-clamp recording under a 60x water immersion objective lens (Olympus, Japan). Neuronal firing and synaptic signals were sampled at 50 kHz using a MultiClamp 700B amplifier (RRID:SCR_018455) and Digidata1550B controlled by pClamp 11 software (Molecular Devices, San Jose, CA; RRID:SCR_011323). Borosilicate glass pipettes (outer diameter = 1.5 mm, inner diameter = 0.86 mm, length = 10 cm, item no. BF150-86-10, Sutter Instruments, Novato, CA) were pulled by micropipette puller (P1000, Sutter instrument, Novato, CA; RRID:SCR_021042) and used for patch clamp recording with a resistance of 3 to 6 MΩ when filled with (i) Cs-methanesulfonate–based internal solution of 120 mM CH_3_O_3_SCs, 2.8 mM NaCl, 10 mM HEPES, 0.1 mM Na_4_-EGTA, 5 mM QX314-HBr, 5 mM phosphocreatine, 0.1 mM spermine, 4 mM ATP-Mg, and 0.4 mM GTP-Na (pH 7.3, 290 mOsm); or (ii) K-gluconate–based internal solution, containing 140 mM K-gluconate, 3.8 mM NaCl, 1 mM MgCl_2_, 10 mM HEPES, 0.1 mM Na_4_-EGTA, 2 mM ATP-Mg, and 0.1 mM GTP-Na (pH 7.3, 290 mOsm); or (iii) high Cl^-^ internal solution, containing: 135 mM CsCl, 3.6 mM NaCl, 10 mM HEPES, 0.1 mM Na_4_-EGTA, 1 MgCl_2_, 10 mM QX314, 2 mM ATP-Mg, and 0.1 mM GTP-Na (pH 7.3, 290 mOsm). Optogenetic stimulation was delivered using a 478-nm light-emitting diode (LED pE-300Ultra, CoolLED, UK; RRID:SCR_021972) through a 60× water immersion objective lens (Olympus), with a field of illumination of ∼450 μm in diameter. Light pulses were delivered to layer 5 or layer 1-3 by slowly moving the objective lens once a whole-cell mode was formed. Peak amplitude of AMPAR-mediated EPSCs or GABA_A_R-mediated IPSCs at −80 mV was quantified as a measure of synaptic strength. The amplitude of NMDA-mediated currents at +40 mV was measured at 50 ms post-optogenetic stimulation when AMPAR EPSCs largely decayed to baseline and used for the measure of NMDA/AMPA ratio. Optogenetically-evoked Sr^2+^-induced asynchronous glutamate release was analyzed between 50 and 500 ms post-optical stimulation. Details of brain slice physiology and optogenetics can be found at Protocols.io (dx.doi.org/10.17504/protocols.io.eq2ly7m2rlx9/v1).

### Behavior

The open field locomotion test was used to assess the development of Parkinsonian motor deficit in MP mice. MP mice and littermates were placed in the center of an open field arena (40 x 40 x 40 cm), and the spontaneous locomotor activity was recorded and analyzed using the Ethovision XT video tracking software (Noldus, Netherlands). The animal was said to have started a bout of movements when it started to move faster than 2 cm/s; and the bout was said to end when the speed of movement dropped below 1 cm/s. Details of the locomotion test can be found at Protocols.io: https://www.protocols.io/view/open-field-locomotion-test-e6nvwjxmdlmk/v1

### Immunohistochemistry

To prepare brain sections for immunohistochemistry, mice were deeply anesthetized using ketamine/xylazine, followed by transcardial perfusion using ice-cold phosphate-buffered saline (PBS; pH = 7.4) for 5 min and paraformaldehyde (PFA, 4%) for 30 min. Brains were then stored in 4% PFA overnight at 4°C before resection. Brain tissues were resected (70 µm in thickness) using a VT1000s vibratome (Leica Biosystems; RRID:SCR_016495). Slices were rinsed three times with PBS before being incubated with 0.5% Triton X-100 and 2% normal donkey serum (Sigma-Aldrich) for 60 min at room temperature, followed by incubation with primary antibodies, including mouse anti-TH (1:2,000; catalog #MAB318, MilliporeSigma; RRID:AB_2201528) or rabbit anti-α5-GABA_A_Rs (1:1000, catalog# PA5-116484, Invitrogen, RRID: AB_2901115) or guinea pig anti-gephyrin (1:1000, catalog# 147318, Synaptic Systems, RRID: AB_2661777), for 48 hours at 4°C or overnight at room temperature. Sections were then rinsed with PBS x3 and incubated with secondary antibodies (1:250) for 90 min, including donkey anti-mouse Alexa Fluor 488 (catalog#715-545-150; Jackson ImmunoResearch Laboratories; RRID:AB_2340846), donkey anti-rabbit Alexa Fluor 594 (catalog# 711-585-152; Jackson ImmunoResearch Laboratories; RRID:AB_2340854), or donkey anti-guinea pig Alexa Fluor 647 (catalog #706-605-148; Jackson ImmunoResearch Laboratories, RRID:AB_2340476), at room temperature before washing with PBS three times. Brain sections were mounted with VectaShield antifade mounting medium (catalog #H-1000, Vector Laboratories; RRID:AB_2336789) and cover-slipped for imaging. TH immunoreactivity (ir) was imaged using a 20X objective lens using a Nikon Confocal microscope (CSU-W1 SoRa). TH immunofluorescence from the dorsal striatum was measured using ImageJ (NIH) and normalized to that from the adjacent cortex to quantify the extent of striatal DA depletion. Details of TH quantification can be found on Protocols.io (https://www.protocols.io/view/quantification-of-th-immunoreactivity-n2bvj85qngk5/v1).

## Confocal microscopy imaging and image analysis

### Dendritic spine analysis

PT and IT neurons were retrogradely labeled using Retrobeads as for electrophysiology studies. Labeled PT and IT neurons were then targeted for whole-cell patch clamp recording using the K-gluconate-based pipette solution containing 2% biocytin. Recorded neurons were filled with biocytin for 40 min and then fixed with 4% PFA for subsequent immunofluorescence and confocal imaging. For dendritic spine analysis of biocytin-filled neurons, segments of apical dendrites from the L2/3 or L5 (20–30 μm in length) were 3D reconstructed using the Filament Tracer function in Imaris (version 9.3, Oxford Instruments, UK; RRID:SCR_007370). The analyzed dendritic segments were located approximately 50–100 μm from the soma. Dendritic spines were manually traced, reconstructed, and quantified for spine density analysis. To quantify dendritic α5-GABARs and gephyrin expression in PT and IT neurons, high-resolution confocal z-stack images were acquired using a Nikon SoRa confocal microscope (Nikon Instruments, Japan) with a 100× oil-immersion objective. Three-dimensional image reconstruction and quantitative analysis were performed using Imaris software (version 9.3, Oxford Instruments, UK; RRID: SCR_007370).

### α5-GABARs and synaptic protein analysis

First, regions of interest (ROIs) encompassing dendritic shafts and spines were manually defined. A 3D isosurface was created for dendritic segments using the Surface function in Imaris. Surface area and volume measurements were obtained from the reconstructed dendritic ROIs. Then, α5-GABAaR or gephyrin immunoreactive puncta were detected using the Spots function in Imaris with a minimal spot diameter of 0.232 μm. Fluorescence thresholds were adjusted to identify receptor puncta while minimizing background noise. Dendritic puncta were determined using the shortest distance-to-surface criterion, with puncta positioned within 0.2 μm of the reconstructed dendritic surface considered dendrite-associated. For co-localization analysis of α5-GABAaRs and gephyrin puncta, dendritic ROIs were first used to generate a masked dataset in Imaris, restricting the analysis to dendritic compartments. Co-localization analysis was then performed using the Coloc module in Imaris.

## Data analysis and statistics

Electrophysiology data were analyzed and quantified using Clampfit 11.1 (Molecular Devices, San Jose, CA; RRID:SCR_011323) and Minianalysis 6.0.3 (Synaptosoft, USA; RRID:SCR_002184). Confocal images were analyzed using ImageJ (NIH) and Imaris (version 9.3, Oxford, UK; RRID:SCR_007370). Statistical analysis was performed in GraphPad Prism 9 (GraphPad Software, San Diego, CA; RRID:SCR_002798). Mann-Whitney U (MWU) test and Wilcoxon signed rank (WSR) test were used for unpaired and paired group comparisons, respectively. Friedman test followed by Dunn’s multiple comparison test or Kruskal-Wallis test followed by Dunn’s multiple comparison test was used to compare means of 3 or more groups. Two-way repeated-measures ANOVA was used to compare group differences in thalamocortical synaptic strength and intrinsic excitability between control and MP mice across a range of stimulation, followed by *post hoc* Sidak’s multiple comparison tests. All statistical tests were two-tailed, and p values of <0.05 (*), 0.01 (**), 0.001 (***), 0.0001 (****) were considered statistically significant. Data were reported as mean and standard error of means (SEM).

## Acknowledgement

We thank the ASAP Team Wichmann for constructive discussion throughout this project. The authors thank Drs. Thomas Wichmann (Emory University) and Stefano Vicini (Georgetown University) for their thoughtful feedback on the manuscript. We thank Dr. Thomas M. Coate (Georgetown University) for providing access to Imaris software and Dr. Larry Zweifel (University of Washington) for sharing CRISPR/SaCas9 AAV vectors. This work was partially supported by research grants from the National Institutes of Health (R01NS121371, H.Y.C.), the Parkinson’s Foundation (PF-IMP-1045313, H.Y.C.), and Aligning Science Across Parkinson’s (ASAP-020572 and ASAP-025187, H.Y.C.) through the Michael J. Fox Foundation for Parkinson’s Research (MJFF). For the purpose of open access, the authors have applied a CC BY public copyright license to all Author Accepted Manuscripts arising from this submission.

## Data and code availability

The raw and analyzed data generated throughout this study have been deposited and are publicly available from: 10.5281/zenodo.21540636.

