## Supplementary figures for "Time Course and Microcircuit Mechanisms of Primary Motor Cortex Dysfunction in Progressive Parkinsonism"

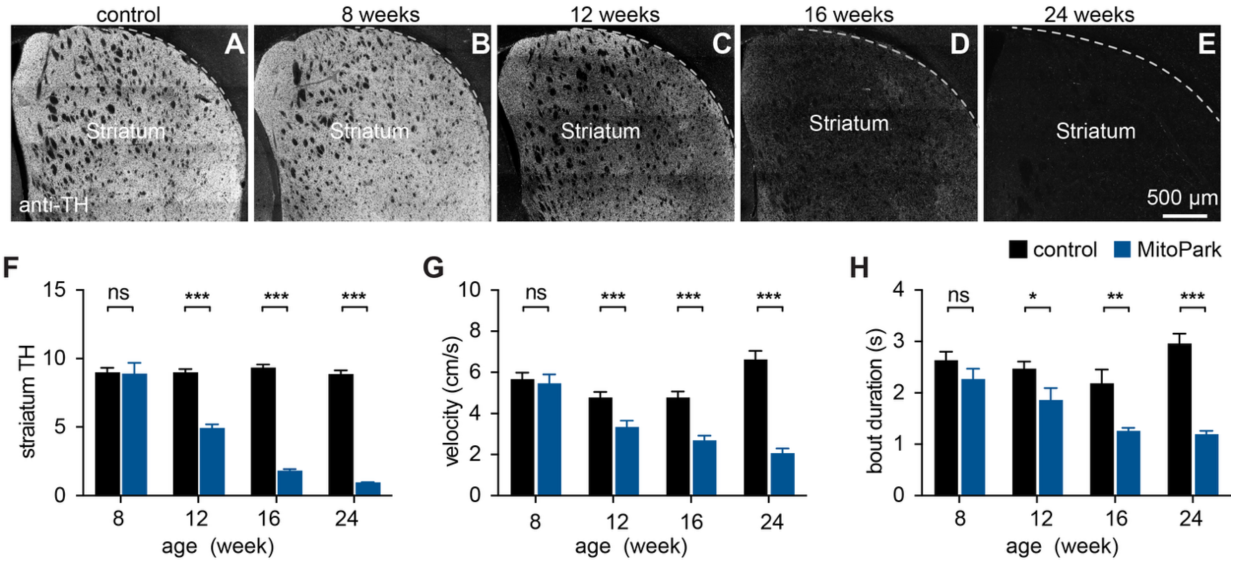


**Figure S1.** MP mice develop age-dependent nigrostriatal degeneration and motor impairments. **A-E**) Representative images showing striatal TH-ir in a 24-week-old control mouse (A) and MP mice across ages (B-E). **F**) Bar graph showing quantification of striatal TH-ir in MP mice at different ages normalized to littermate controls. At 8 weeks, N = 5 controls and 4 MP mice; at 12 weeks, N = 10 controls and 10 MP mice; at 16 weeks, N = 9 controls and 9 MP mice; at 24 weeks, N = 12 controls and 14 MP mice. To determine striatal TH levels, TH immunofluorescence in the dorsal striatum was normalized to the adjacent cortex. **G-H**) MP mice develop progressive reduction in the speed of locomotor activity (G) and duration of mobile bouts (H). At 8 weeks, N = 8 controls and 7 MP mice; at 12 weeks, N = 12 controls and 12 MP mice; at 16 weeks, N = 8 controls and 8 MP mice; at 24 weeks, N = 8 controls and 8 MP mice. Two-way ANOVA followed by Sidak test. ns, not significant. **p < 0.01, ***p < 0.001.

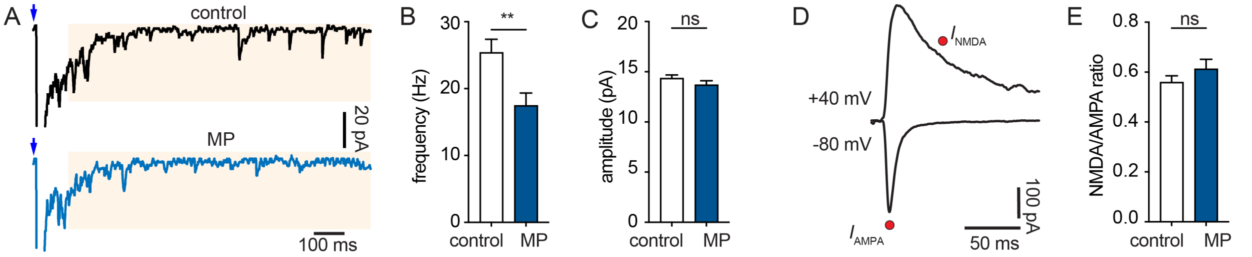


**Figure S2. Decreased functional thalamo-PT connections in 24-week-old MP mice**. **A**) Representative traces showing optogenetically-evoked asynchronous EPSCs in PT neurons from control and MP mice at 24 weeks of age. Blue arrows indicate optogenetic stimulation. Yellow shaded areas indicate the time window for measuring asynchronous EPSCs. **B-C**) Summary graph showing the frequency and amplitude of asynchronous EPSCs at the thalamo-PT synapses. Frequency: control = 25.4±1.98 Hz, MP = 17.5±1.88 Hz, p = 0.003. Amplitude: control = 14.4±0.33 pA, MP = 13.7±0.41 pA, p = 0.13. Control = 30 cells/3 mice; MP = 38 cells/4 mice, MWU test. **D**) Representative traces of EPSCs at -70 mV and +40 mV. Red dots indicate where AMPAR- and NMDAR-EPSCs were measured. **E**) Summary graph showing NMDA/AMPA ratio between control and MP mice at 24 weeks of age. control = 0.56±0.026, n = 21 cells/3 mice, MP = 0.061±0.039, n= 27 cells/ 4 mice. p = 0.37, MWU test.

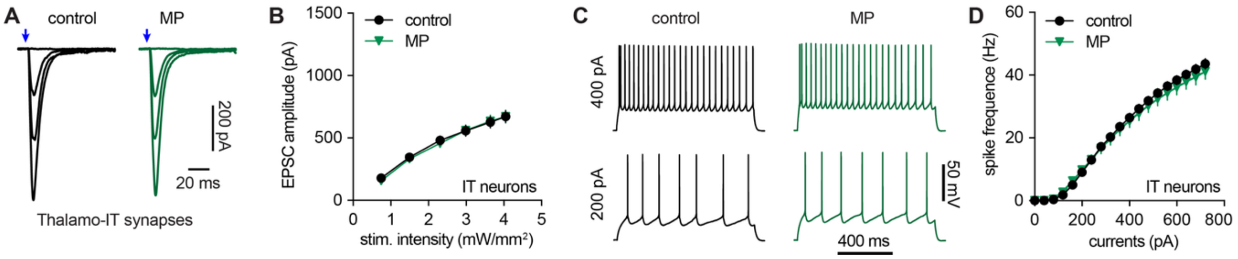


**Figure S3. Physiological properties of IT neurons in M1 are not altered in advanced MP mice. A-B**) Synaptic strength of thalamic inputs to IT neurons is unaltered in 24-week-old MP mice (n = 30 neurons/4 mice) compared to littermate controls (n = 31 neurons/4 mice). **C-D**) Intrinsic excitability of IT neurons in M1 is not altered in 24-week-old MP mice compared to littermate controls. Control = 32 neurons/6 mice, MP = 34 neurons/6 mice. Two-way RM ANOVA for statistical comparisons in B and D.

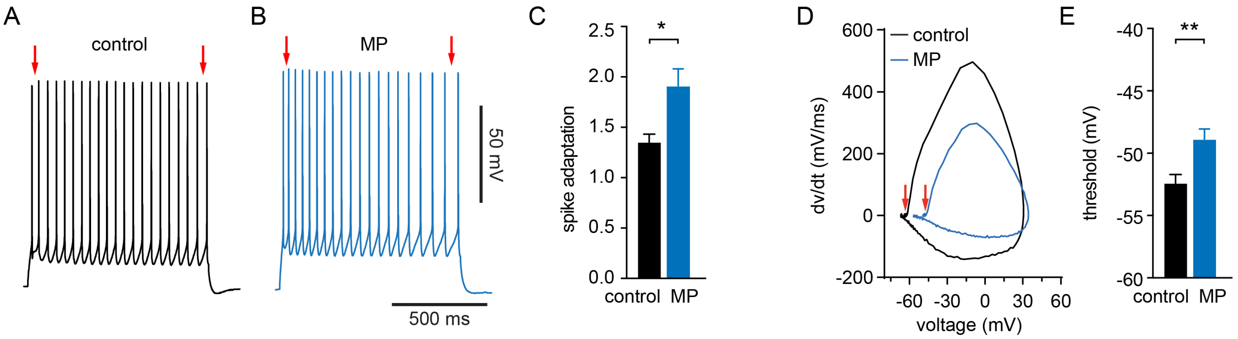


**Figure S4. Physiological properties related to altered intrinsic excitability of PT neurons in 24-week-old MP mice. A-C)** PT neurons from 24-week-old MP mice show stronger spike adaptation during repetitive firing than those from littermate controls. Red arrows in A and B indicate the first and last inter-spike intervals used for calculating spike adaptation index. Spike adaptative index, control = 1.35±0.08, n = 50 neurons/10 mice, MP = 1.91±0.17, n = 33 neurons/8 mice; p = 0.02, Mann-Whitney U test. **D-F**) Representative AP phase plane plots showing depolarized AP threshold of PT neurons from MP mice compared to those of littermate controls at 24 weeks of age. Threshold (red arrows), control = -52.4±0.72 mV, n = 50 neurons/10 mice, MP = -48.9±0.83 mV, n = 33 neurons/8 mice; p = 0.002, MWU.

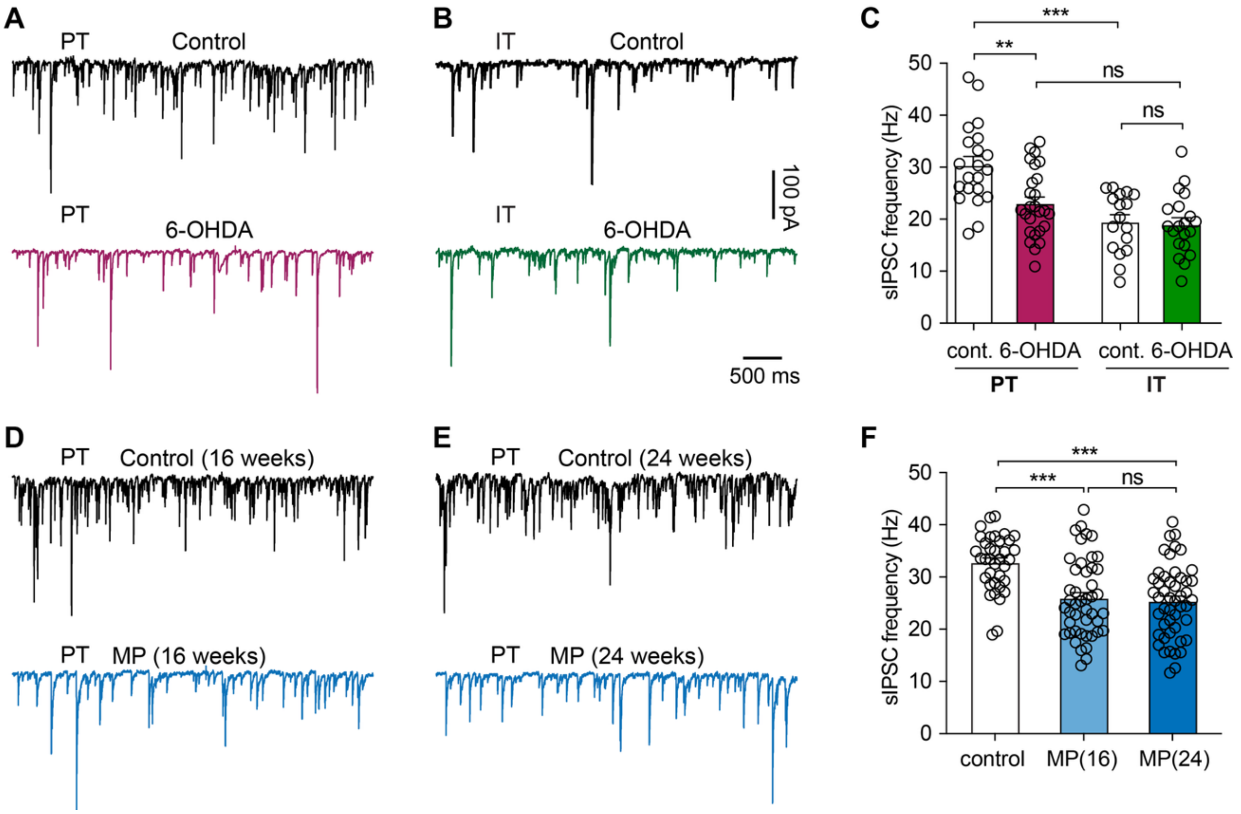


**Figure S5. Subtype-specific inhibition of M1 pyramidal neurons. A-B)** Representative sIPSC traces of PT (A) and IT (B) neurons from control and 6-OHDA-lesioned mice using a high $Cl^-$ internal solution and in the presence of DNQX (10 μM) and D-APV (50 μM) in the bath solution to block ionotropic glutamatergic transmission. **C**) Bar graph showing the difference in the sIPSC frequency of PT and IT neurons under healthy conditions, which was abolished by loss of SNc DA neurons in the parkinsonian state. sIPSC frequency of PT neurons, control = 30.4±1.7 Hz, n = 21 neurons/3 mice; 6-OHDA = 23.0±1.3 Hz, n = 26 neurons/4 mice; p = 0.0017; sIPSC frequency of IT neurons, control = 19.5±1.4 Hz, n = 17 neurons/3 mice; 6-OHDA mice = 19.0±1.3 Hz, n = 20 neurons/4 mice; p > 0.99; Two-way ANOVA followed by Sidak test. **D-E**) Representative sIPSC traces of PT neurons from control and MP mice at 16 (D) and 24 (E) weeks old. **F**) Summary graph showing the decreased sIPSC frequency of PT neurons during the gradual loss of SNc DA neurons in the MP mice. sIPSC frequency, control = 32.8±0.9 Hz, n = 36 neurons/4 mice; MP at 16 weeks = 25.9±1.1 Hz, n = 45 neurons/4 mice; MP at 24 weeks = 25.4±1.0 Hz, n = 48 neurons/4 mice. Data from control mice at 16- and 24-weeks of age were combined in F. ns, not significant; ***, p < 0.001; Kruskal-Wallis test followed by Dunn tests.

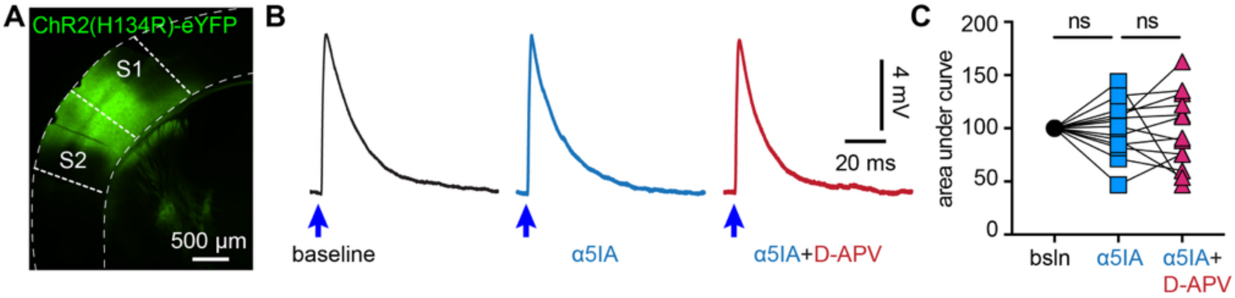


**Figure S6. α5-GABA$_A$Rs do not modulate corticocortical transmission. A**). Representative AAV-ChR2 infection area in the sensory cortices. **B**). Representative PSPs that were evoked by optogenetic stimulation of sensory cortical inputs to M1 PT neurons at baseline, in the presence of α5IA, and after the application of α5IA plus D-APV. **C**). Summarized results showing the effects of α5IA and α5IA plus D-APV on the area under the curve of PSP at baseline, in the presence of α5IA (% baseline = 99.31±7.06%), and after the application of α5IA plus D-APV (% baseline, 97.62±9.76%). n = 13 neurons/3 mice. ns. Not significant. Friedman test.

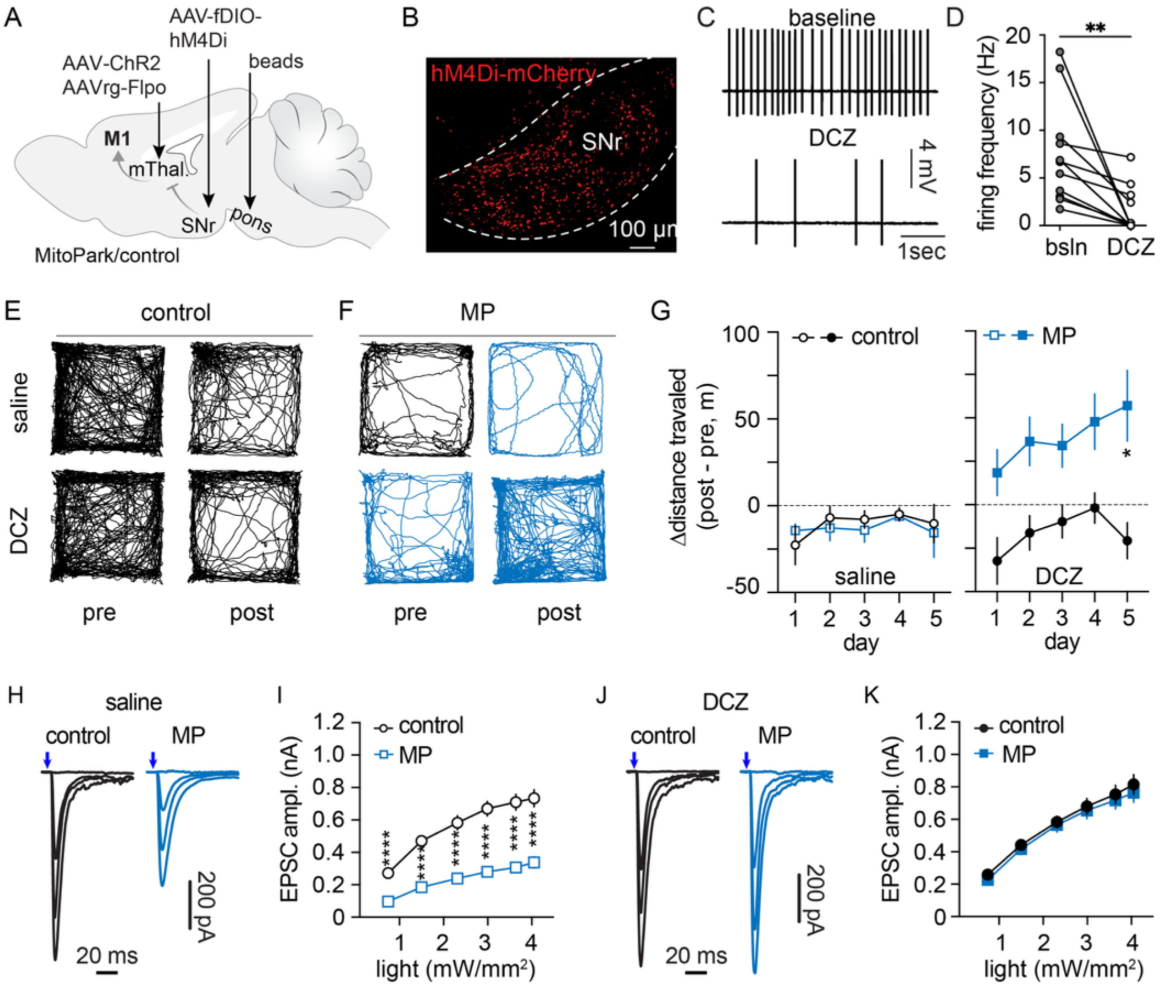


**Figure S7. Chemogenetic suppression of basal ganglia output restores thalamo-PT synaptic strength in advanced MP mice. A**) Experimental design. **B**) Confocal image showing hM4Di-mCherry-expressing cells in the SNc. **C-D**) Selective DREADs agonist DCZ suppresses the autonomous firing of SNr neurons in brain slices. Firing rate, baseline = 7.55±1.6, DCZ = 1.59±0.74, n =11 cells/5 mice, p = 0.001, Wilcoxon test. **E-F**) Representative track plots showing locomotor activities of control (E) and MP mice (F) in response to saline (top) and DCZ (bottom) injections. **G**) Summary data showing changes in locomotor activity of control and MP mice after saline (left) or DCZ (right) injections relative to baseline. Two-Way RM ANOVA followed by Sidak tests. **H-I**) Representative traces showing the thalamo-PT synaptic transmission in saline-injected control and MP mice. **J-K**) Representative traces showing the thalamo-PT synaptic transmission in DCZ-injected control and MP mice. Two-way RM ANOVA followed by Sidak tests. *p < 0.05, **p < 0.01, ***p < 0.001, ****p < 0.0001.
