## Supplementary table 1 for "Time Course and Microcircuit Mechanisms of Primary Motor Cortex Dysfunction in Progressive Parkinsonism"

### Key resources table

| REAGENT OR RESOURCE | SOURCE | IDENTIFIER |
|---|---|---|
| **Antibodies** | | |
| Anti-mouse tyrosine hydroxylase | MilliporeSigma | RRID: AB_2201528 |
| Anti-Rabbit GABA-A-α5 | Thermo Fisher Scientific | RRID: AB_2901115 |
| Anti-Guinea pig Gephyrin | Synaptic System | RRID: AB_2661777 |
| Alexa Fluor 647 donkey anti-mouse IgG | Jackson ImmunoResearch Labs | RRID: AB_2340862 |
| Alexa Fluor 488 donkey anti-mouse IgG | Jackson ImmunoResearch Labs | RRID: AB_2341099 |
| Alexa Fluor 594 donkey anti-mouse IgG | Jackson ImmunoResearch Labs | RRID: AB_2340858 |
| **Chemicals and Virus** | | |
| AAV9-hSyn-ChR2 (H134R)-eYFP | Addgene | RRID: Addgene_26973 |
| pAAV-EF1a-Flpo-AAV Retrograde | Addgene | RRID: Addgene_55637 |
| AAV9-FLEXfrt-SaCas9-U6-sgGrin2b | Vector biolabs | RRID: Vector biolabs_25110#53 |
| AAV1-EF1α-fDIO-tdTomato | Addgene | RRID: Addgene_128434 |
| AAV1-FLEXfrt-sgRosa26 | Gift from Larry Zweifel (University of Washington) | n/a |
| Red retrobeads | Lumafluor Inc. | n/a |
| Fast blue | Polyscience | RRID: Polyscience_17740-1 |
| Tetrodotoxin (TTX) | Hellobio. | HB1034 |
| 4-Aminopyridine (4-AP) | Hellobio. | HB1073 |
| SR95531 hydrobromide (Gabazine) | Hellobio. | HB0901 |

| | | |
|---|---|---|
| D-AP5 | Hellobio. | HB0225 |
| DNQX disodium salt | Hellobio. | HB0262 |
| Ifenprodil | Hellobio. | HB0339 |
| L-DOPA | Hellobio. | HB1925 |
| Desipramine hydrochloride | MilliporeSigma | D3900 |
| Pargyline hydrochloride | MilliporeSigma | P8013 |
| 6-Hydroxydopamine (6-OHDA) hydrobromide | Hellobio. | HB1889 |
| α5IA | Santa Cruz Biotechnology | sc-252345 |
| NaHCO3 | Fisher Chemical | S369 |
| KCl | Fisher Chemical | P217 |
| $NaH_2PO_4H_2O$ | Fisher Chemical | S369 |
| $CaCl_2.2H_2O$ | Fisher Chemical | C79 |
| $MgSO_4.7H_2O$ | Fisher Chemical | M63 |
| D-Glucose | Fisher Chemical | D16 |
| Sucrose | Fisher Chemical | S5 |
| HEPES | Millopore-Sigma | H3375 |
| Phosphocreatine | Millopore-Sigma | P1937 |
| $Na_4EGTA$ | Millopore-Sigma | E8145 |
| $Na_3GTP$ | Millopore-Sigma | 51120 |
| $Mg_{1.5}ATP$ | Millopore-Sigma | A9187 |
| K-Gluconate | Tokyo Chemical Industry | G0040 |
| TEA-Cl | Millopore-Sigma | T2265 |
| QX314 HBr | Hellobio. | HB1029 |
| Spermine | Millopore-Sigma | S3256 |
| CsCl | Millopore-Sigma | 289329 |
| CsOH | Millopore-Sigma | C8518 |

| | | |
|---|---|---|
| Strontium Cl | ThermoFisher Scientific | 369740250 |
| L-Ascorbic acid | Millopore-Sigma | A5960 |
| Desipramine hydrochloride | Millopore-Sigma | D3900 |
| Pargyline hydrochloride | Millopore-Sigma | P8013 |
| **Animals** | | |
| $DAT^{Cre}$ mice | Jackson Labs | RRID: JAX_006660 |
| $Tfam^{fl/fl}$ mice | Jackson Labs | RRID: JAX_026123 |
| $SST^{Cre}$ mice | Jackson Labs | RRID: JAX_013044 |
| Ai32 reporter mice | Jackson Labs | RRID: JAX_024109 |
| C57BL/6 | Charis River | RRID:MGI:2159769 |
| **Software** | | |
| ImageJ | NIH | https://imagej.nih.gov/ij/ |
| Prism | Graphpad | https://www.graphpad.com/ |
| Clampfit | Molecular Devices | n/a |
| Adobe illustrator | Adobe | n/a |
| EthoVision XT 17 | EthoVision | n/a |
| Imaris | Oxford Instruments | https://imaris.oxinst.com/versions/10-1 |
| Multiclamp 700B | Molecular Devices | n/a |
| Digidata 1550B | Molecular Devices | n/a |
